# Comprehensive Multi-omics Mapping of Vesicle Cargo from Plasma or Novel Tissue Vesicles Reveals Pathological Changes in Cargo with Patient Age

**DOI:** 10.1101/2024.12.06.627231

**Authors:** George Ronan, Lauren Hawthorne, Jun Yang, Ruyu Zhou, Fang Liu, Pinar Zorlutuna

## Abstract

Aging is a major risk factor for cardiovascular disease, the leading cause of death worldwide, and numerous other diseases, but the mechanisms of these aging-related effects remain elusive. Recent evidence suggests that chronic changes in the microenvironment and local paracrine signaling are major drivers of these effects, but the precise effect of aging on these factors remains understudied. Here, for the first time, we directly compare extracellular vesicles obtained from young and aged patients to identify therapeutic or disease-associated agents, and directly compare vesicles isolated from heart tissue matrix (TEVs) or plasma (PEVs). While young TEVs and PEVs showed notable overlap of miRNA cargo, aged EVs differed substantially, indicating differential aging-related changes between TEVs and PEVs. TEVs overall were uniquely enriched in miRNAs which directly or indirectly demonstrate cardioprotective effects, with 45 potential therapeutic agents identified in our analysis. Both populations also showed increased predisposition to disease with aging, though through different mechanisms. Changes in PEV cargo were largely correlated with chronic systemic inflammation, while those in TEVs were more related to cardiac homeostasis and local inflammation. From this, 17 protein targets were identified which were unique to TEVs and highly correlated with aging and the onset of cardiovascular disease. Further analysis via machine learning techniques implicated several new miRNA and protein targets, independently suggesting several of the targets identified by non-machine learning analysis, which correlated with aging-related changes in TEVs. With further study, this biomarker set may serve as a powerful, potential indicator of cardiac health and age which can be measured from PEVs. Additionally, several proposed “young-enriched” therapeutic agents were validated and, when tested, could successfully prevent cell death and cardiac fibrosis in disease-like conditions.

## INTRODUCTION

Advanced age is quickly becoming the most prevalent risk factor worldwide for numerous diseases, including cardiovascular disease (CVD), leading cause of death worldwide^1^, obesity and obesity-related diseases^2^, and many cancers^3^, and by 2050 the global population aged 65 or older is expected to double to over 2 billion or nearly 20% of predicted global population^4^. In some rapidly aging countries, such as Japan and Italy, 65+ aged individuals already account for over 20% of the population^5^, with Japan quickly approaching 30%. Recent literature has implicated long-term changes in the cardiac microenvironment in aging-related CVD pathogenesis^6–9^, and suggests that these changes can be manipulated to improve patient outcomes. The use of extracellular matrix (ECM) or ECM-derived materials has seen some success in pre-clinical trials by leveraging previously established effects of ECM treatment, including immunomodulation^7,10,11^, stem cell recruitment^7,12,13^, and functional revitalization^6,12–14^. These properties enhance the reparative effects of novel biomaterials, commonly being integrated with hydrogels or other bioscaffolds in treatment of CVDs^9,10,13,15,16^. However, the mechanisms by which ECM promotes these effects is not well understood, and the drivers of aging-related changes in the microenvironment are critically understudied.

Extracellular vesicles (EVs) have been recently identified as potential agents of aging-related changes in the microenvironment^6,7,14^ and key mediators of many therapeutic effects observed in ECM treatment^11,13,17^. However, many of these studies have been largely limited to EVs isolated from plasma and have been identified as exosomes^18,19^. Exosomes are a specific subcategory of small EV (sEV), or EVs with typical sizes of under 200 nm^20,21^, that express characteristic surface protein markers and serve as common vehicles for endocrine, paracrine, and even autocrine transport of RNAs and proteins^21–23^. While more recent efforts have focused on the broader category of sEVs, due to difficulties in separating exosomes from other sEV subgroups such as supermeres and exomeres^7,24^, exosomes have been investigated as “tissue-free” theragnostic nanoparticles. Stem cell-derived exosomes are being investigated in a clinical trial as vital components for an “off-the-shelf” cardiac patch that emulates the benefits of direct ECM incorporation^13,25^, and plasma exosomes have provided a wealth of real-time diagnostic information for patients in both CVDs^15,26^ and cancer diagnosis^27,28^.

Tissue matrix-derived sEVs provide a unique avenue to evaluate the effects of aging-related changes in the microenvironment on the progression of CVDs, a topic that has remained elusive and difficult to address clinically^8,10,13,14,17,29^. While CVD-related mortality is often actuated by the onset of myocardial infarction (MI)^1,30^, the risk of mortality is predominantly regulated by a combination of risk factors including previous cardiac event^30–32^, obesity and metabolic health^30,33^, and age^34,35^, and recent studies have clearly demonstrated correlative effects between age and the risk of MI or chronic CVD^4,8,35–37^.

Previously, we discovered and characterized cardiac tissue ECM-resident exosome-like sEVs from donor hearts from young or aged donors, and found that the size, morphology, and miRNA and protein contents of these sEVs significantly changed with donor age^7^. Despite a growing body of literature supporting the use of sEVs as dynamic theragnostic tools in CVD and other diseases^9,17^, currently in literature there is no single resource that directly investigates and establishes aging-related changes in sEV content for tissue-resident sEVs, and any head-to-head comparison of miRNA and whole protein profiles of tissue-resident or plasma-derived sEVs is similarly absent. This may be due to the relatively recent discovery of ECM-resident sEVs^38^, but remains a notable gap in knowledge.

Herein we, for the first time in literature, characterize differences in size profile, surface protein distribution, and uptake mechanics of sEVs derived from cardiac tissue ECM, or tissue sEVs (TEVs), and sEVs derived from plasma, or plasma sEVs (PEVs). We also establish previously unreported differential aging-related changes in these metrics between TEVs and PEVs by assessing both populations of EVs from young (<40 y.o., n = 6 for tissue & plasma) and aged (>50 y.o., n = 6 for tissue & plasma) donors. After this, we provide a whole miRNA profile for TEVs and PEVs from young and aged donors, a dataset currently absent in literature, and identify targets and pathways of interest for CVD within our dataset. Following this, we do the same for a whole protein profile, again from TEVs and PEVs from both young and aged donors for the first time in literature. From these analyses, we can successfully bifurcate the cargo profiles of TEVs and PEVs, with PEVs tending to be largely correlated with systemic immune response and cardiac TEVs tending to be related to cardiac homeostasis and CVD, as expected. TEVs were also uniquely enriched in cardioprotective miRNAs involved in the regulation and prevention of fibrosis and anoxia-related cell death. Most interestingly, aged-source EVs in both populations tended to exhibit higher enrichment of cargo involved in disease-associated pathways than young-source EVs not through nucleic acid cargo, but through chauffeured proteins. While the specific pathways involved were different, with aged PEVs mostly trafficking proteins related to increases in chronic systemic inflammation and neurological diseases and aged TEVs transporting proteins relating to increased risk of heart failure, increased local inflammation, and overall metabolic shift, both populations suggested that EVs can propagate aging-related damage-inducing proteins. From these cargo analyses, we identified 45 miRNA targets which provide cardioprotective effects and 17 proteins which are implicated in the development of CVDs. Additionally, we utilize machine learning algorithms to identify biomarkers of cardiac aging which can be detected from PEVs, providing a quantitative, low-invasive method of assessing long-term cardiac health or heart “age”. In addition to the identification of biomarkers, this analysis also validated several identified potential therapeutic miRNAs. These miRNAs, in combination with previously evaluated therapeutic miRNAs, were applied to cells in an MI-like environment to improve recovery post-MI, and successfully improved survival of both muscle and stromal cells.

With these data we achieve several goals. First, we provide a resource for future studies investigating aging-related changes in tissue-resident sEVs, a recently discovered source of EVs that are currently poorly understood but provide exciting avenues for understanding and manipulating disease-related intra-tissue signaling. Second, we contrast both these tissue-resident EVs and classical plasma EVs as well as directly assess aged and young cargo from both populations to identify aging-related changes in EV content and signaling, a sorely understudied topic in EV research. Third, from the EV cargo we have identified both therapeutic miRNAs and damage-associated protein cargo to direct future target-based studies and drug development for notoriously difficult to target diseases, such as cardiac fibrosis. The dissemination of these data will help promote more in-depth study of sEVs as both diagnostic and therapeutic vehicles to investigate the effects of chronic changes to tissues, such as aging, and provide a useful tool for the future investigation of comprehensive comparison of sEV populations. Fourth, we identified both miRNA and protein biomarkers which, in TEVs, were indicative of aging and related to CVD onset, but could also be identified from PEVs of aged donors. These biomarkers suggest that, with greater understanding of aging-related changes in PEV and TEV cargo, it will be possible to identify quantitative biomarkers of tissue aging from PEVs, as opposed to biopsy or similarly invasive techniques. Lastly, we validated our previous hypothesis that further comparison of TEVs and PEVs from differentially aged cohorts would identify further therapeutic miRNAs^7^. While this therapeutic demands further testing, the therapeutic efficacy of this cocktail in a simple lipid nanoparticle delivery mechanism compared to TEVs directly is an indicator that the therapeutic effects of TEVs can in fact be reconstructed from specific, young-enriched cargos.

## RESULTS

### Characteristics of sEV size profile change depending on biological source

Transmission electron microscopy (TEM) of sEVs isolated from tissue (TEVs) or plasma (PEVs) of young (<40 y.o., n = 6 for tissue & plasma, Supplemental Table S1) and aged (>50 y.o., n = 6 for tissue & plasma, Supplemental Table S1) donors showed that all particles were similar in overall morphology and staining characteristics (Figure 1A). sEV size profile was validated via Nanoparticle Tracking Analysis (NTA), and the resultant statistics showed that size was mostly consistent between young and aged samples in PEVs (young mode: 129.5 nm; aged mode: 159.5 nm) but not TEVs (young mode: 128.5 nm; aged mode: 49.5 nm) (Figure 1B). Furthermore, TEVs were predominantly smaller than PEVs overall, although all sEVs fell within the expected size range for exosome-like sEVs. All populations were found to be monodisperse, though young tissue-source TEVs had a notably larger PDI than any other cohort (Supplemental Table S2).

**Figure 1.**
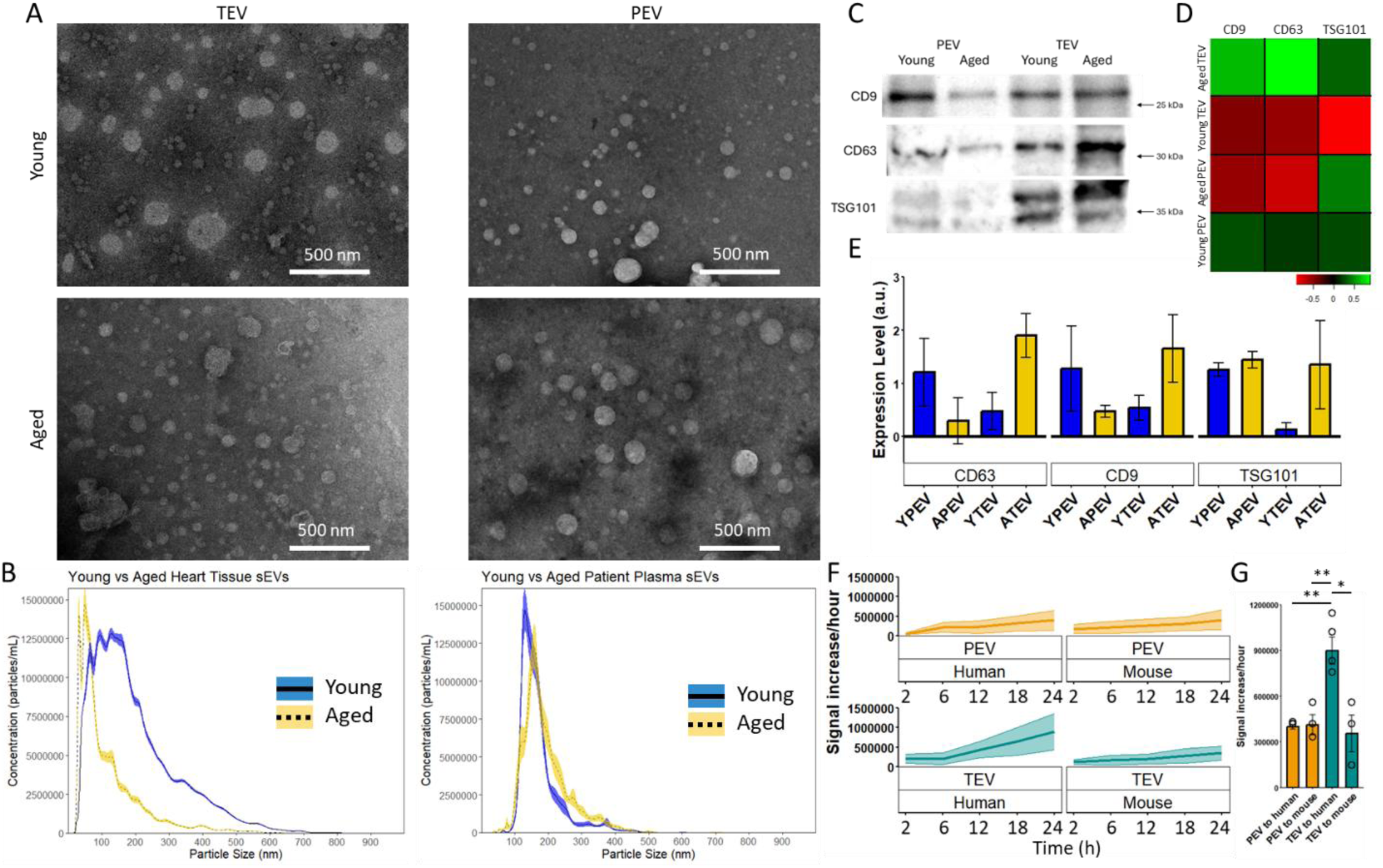
Tissue resident sEVs display substantive differences from those isolated from plasma in size profile, surface protein expression, and behavior. Measurements were taken from n = 6 biological replicates per cohort, equally spread between male and female donors. (A) Transmission electron microscopy representative images for each cohort. (B) Nanoparticle tracking analysis differentiating young (blue) or aged (gold) source tissue sEVs (left) or plasma sEVs (right). The overall expression of exosomal surface proteins was measured by western blot (C) and compared (D) and the change in expression between groups was compared with respect to tetraspanin (E). (F) Uptake rate of TEVs or PEVs from young donors (n >= 3 for each) on human or mouse fibroblasts in 2D culture (n >= 3 replicates for each) over 24 hours, presented as change in uptake rate per hour between timepoints taken ± standard error. (G) Endpoint uptake rate of TEVs or PEVs on human or mouse fibroblasts, bars represent mean of biological replicates ± standard deviation and points represent mean of replicates for each biological replicate. * p < 0.05, ** p < 0.01, assessed by Welch’s t-test for (E) and (G).

### Different tissue source sEVs demonstrate differential responses to aging effects

Western blot was performed to identify characteristic exosome markers CD9, CD63, and TSG101 on the sEVs isolated, with each protein showing up clearly (Figure 1C). The measured quantity of each protein was also assessed from the blot using ImageJ for standardized blot quantification, and the results were Z-scored for comparison (Figure 1D). These show that changes in expression of these particular exosomal surface proteins behave inversely with respect to age between TEVs and PEVs, with PEVs showing a 0.5-fold to 1-fold decrease in expression of CD9 and CD63 between the young and aged cohorts and TEVs showing a greater than 1-fold increase in expression across that same comparison (Figure 1E). CD63 expression remained mostly consistent with CD9 expression in both populations with respect to age, though the change in expression was opposite between TEVs and PEVs, but TSG101 availability was increased with age only in TEVs while remaining consistent in PEVs, although it should be noted that these differences were not significant with the metrics used. Taken with the above results, this suggests a shift in the tetraspanin web and targeting abilities^39–41^ that substantially differs between the TEVs and PEVs for the proteins assessed.

### Uptake kinetics of sEVs varies between tissue source and destination tissue

Uptake kinetics of sEVs with typical or atypical cell culture populations was also assessed (Figure 1F). When applied to murine cells, both PEVs and TEVs maintained steady uptake by cells over 24 h, resulting in a nearly flat line (right). When applied to human cardiac cells, however, PEVs showed a slight increase in uptake over time whereas TEVs showed exponential increase in uptake rate over 24 h, reaching several-fold higher uptake rates and uptake overall than PEVs (Figure 1G). Additional assays with young vs aged cells (artificially aged iCFs or aged mice) showed that the age of the cell had no significant effect on either the uptake rate of EVs or the size of the EVs produced (Supplemental Figure S1), an unexpected result.

### Profiling of miRNA content

miRNA profiling via Nanostring analysis revealed highly distinct miRNA profiles between PEVs and TEVs (Figure 2A), with sources tending to cluster together both when compiled and in individual replicates (Supplemental Figure S2). While sEVs from young donors of both sources showed some notable overlap, the miRNA profiles of sEVs from aged donors were almost completely distinct, with less than 100 miRNAs in common. Relative expression values for all 798 detected miRNA expressed in these samples is provided in the supplement (Supplemental Table S3). Prevalence of miRNAs in all sEVs was mostly normally distributed (Figure 2B), though both PEV populations had a notable number of very highly expressed (detected count >30 in chip) miRNAs. In general, sEVs from young donors tended to express a larger number of miRNAs overall but at lower counts, while sEVs from aged donors tended to express fewer miRNAs but at higher concentrations, consistent with previous findings^7^. Unsupervised analysis of the data to identify distinguishing miRNAs between TEVs and PEVs revealed 125 potential targets (Figure 2C). Of these 125 miRNAs, 59 were expressed in TEVs but not PEVs, and 66 were highly expressed in PEVs but not TEVs (Supplemental Table S4).

**Figure 2.**
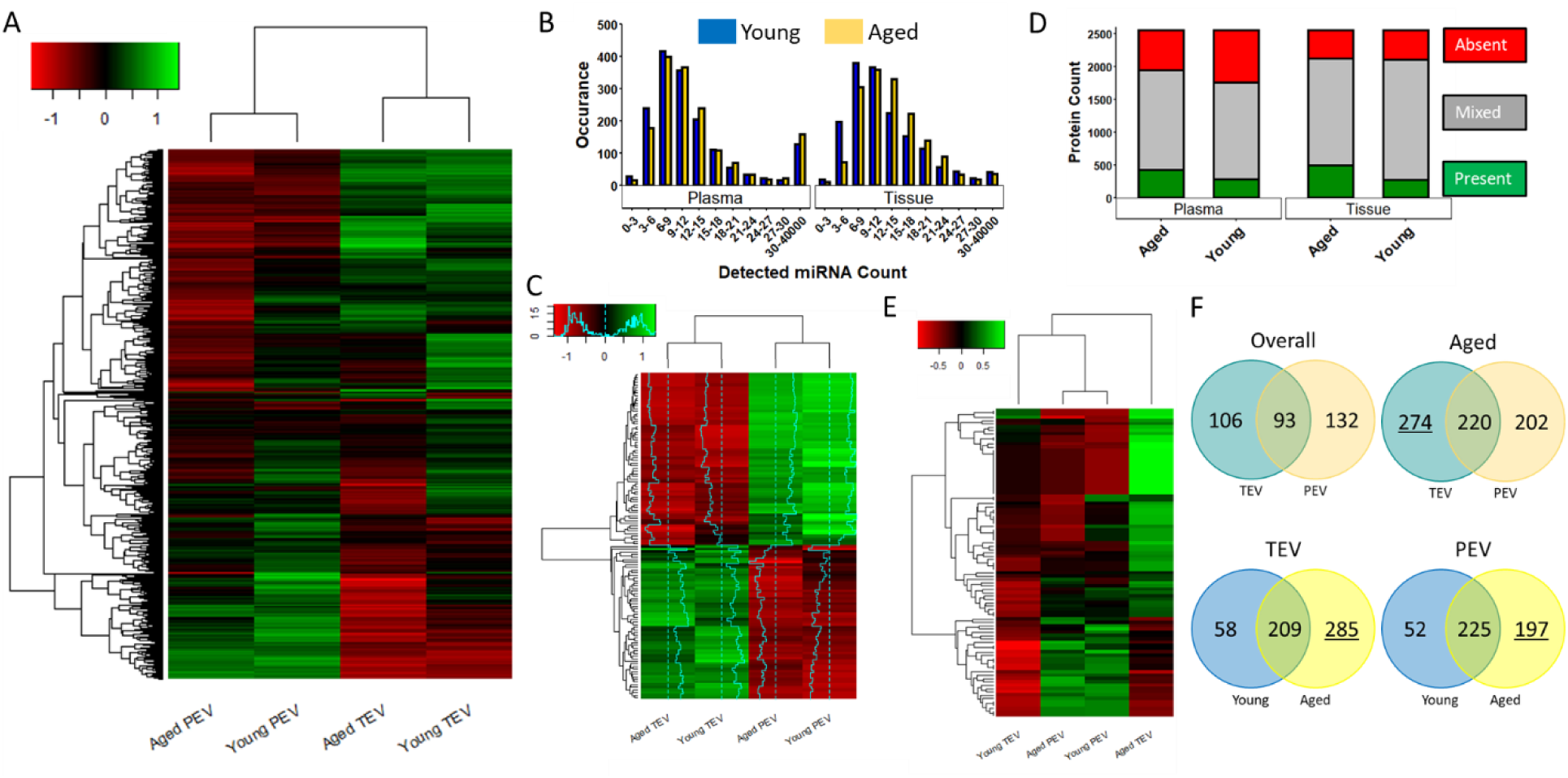
Summary of total miRNA and protein profile from tissue or plasma-source sEVs. Measurements were taken from n = 6 biological replicates per cohort, equally spread between male and female donors. (A) Heatmap showing the full miRNA profile for each cohort. Supplementary figures are available for individual replicates (Supplemental Figure S2) and for separation by both age and sex (Supplemental Figure S3) of total miRNA profile. (B) Histogram detailing the overall prevalence and distribution of miRNAs measured for each cohort. (C) Heatmap of the miRNA targets identified via unsupervised analysis that distinguish between TEVs and PEVs. A total of 125 miRNAs were identified and were nearly evenly split between TEVs and PEVs, with TEVs overexpressing 59 miRNAs and PEVs claiming 66. A full list of these miRNAs can be found in the supplement (Supplemental Table S4). (D) Stacked bar plot showing the expression profile of all 2547 detected proteins for individual cohorts, separating between proteins that are absent in all biological replicates (red, top), present in all biological replicates (green, bottom), or neither (grey, middle). (E) Heatmap showing the relative expression of the 93 proteins that were expressed in every biological replicate across all cohorts (Supplemental Table S7). Supplementary figures are available of all 2547 detected proteins for cohorts and individual replicates (Supplemental Figure S7) and for separation by both age and sex (Supplemental Figure S8) of both protein profiles. (F) Venn diagrams showing overlap of “present” proteins across cohorts, comparing between TEVs and PEVs overall (top left), only aged donor TEVs and PEVs (top right), young and aged donor TEVs (bottom left), and young and aged PEVs (bottom right). In comparisons where one side shows >33% greater quantity of proteins, the number of proteins was underlined.) * p < 0.05.

### Profiling of protein content

Protein profiling via mass spectrometry and subsequent analysis revealed that TEVs and PEVs tended to separate into two distinct clusters depending on tissue source, although this was less consistent when comparing individual biological replicates (Supplemental Figure S7). This may be due to some proteins being entirely absent from detection in many biological replicates (Figure 2D). Visualization of the overall protein population in terms of prevalence or absence of proteins by number also revealed that many proteins were enriched inconsistently between biological replicates, with less than 500 (< 20% of all identified proteins via mass spec) were universally present in any given cohort, and a similar number was universally absent in every cohort. To present a more readily comparable visual, the 93 proteins common across all biological replicates for all cohorts were also mapped (Figure 2E, Supplemental Figure S7, Supplemental Table S7). While in the overall population TEVs showed a slightly higher overall protein enrichment, with aged donor TEVs still having higher overall protein enrichment than young donor TEVs, comparison of only the common proteins across all replicates revealed a substantial increase in enrichment in aged donor TEVs compared to all groups while young donor TEVs displayed notably low levels of relative protein enrichment. Further subdivision of cohorts by sex in addition to age additionally reveals that this is not a sex-dependent shift, though the overall protein enrichment may be sex-dependent (Supplemental Figure S8). This is consistent with our previous findings on relative protein content between aged and young donor TEVs^7^.

### Aging-correlated shifts in protein profile is similar between TEVs and PEVs

Breakdown of protein enrichment by cohort was recorded via Venn Diagram (Figure 2F), and reveals that overall both TEVs and PEVs have a similar number of overlapping and distinctive protein cargo. Both populations, however, show similar changes in cargo with respect to age. In both TEVs and PEVs, comparison of young donor sEV unique protein contents with that of aged donor sEV content reveals a nearly 3-fold increase in unique proteins consistently expressed in all biological replicates. Aged donor TEVs have a higher number of unique proteins compared to TEVs.

### Mapping of miRNA targets implicates TEV signaling in multi-point regulation of cardiovascular health-related pathways

Of the 59 miRNA targets overexpressed in TEVs, 45 have been identified as “potentially involved” in CVD according to PubMed and other online databases^26^. These 45 miRNAs were pulled out and visualized (Figure 3A), evaluated for potential gene targets and overlap (Figure 3B, C), assessed via gene ontology (Figure 3D), and mapped using MetaCore Pathway Analysis (Figure 3E-H). Likely gene targets were assessed for overlapping targets of miRNAs and contributions of miRNAs to individual gene targets. Analysis revealed 11,366 potential gene targets (Supplemental Table S5), though more than two-thirds were targeted by only a single target miRNA. Of the target miRNAs, miR-519, miR-181, and miR-548 contributed to the largest number of gene targets (Supplemental Figure S4). Genes which were targeted by 10 or more target miRNAs were selected for gene ontology analysis, resulting in selection of 167 genes (Supplemental Figure S4, Supplemental Table S6). Within this cohort of genes, miR-548 and miR-519 remained the top contributors with miR-129 supplanting miR-181, though miR-181 remained a major contributor (Figure 3B). The top 10 targeted genes were also identified (Supplemental Figure S5) along with contributing miRNAs (Figure 3C, Supplemental Figure S5). Of the top 10, the disruption of regular signaling for ZBTB20, CLOCK, TNRC6B, RORA, CREB1, MECP2, MINDY2, TAOK1, and AAK1 was linked to cardiovascular disease^42–51^, typically through analogous pathways to chronic CVD or similar chronic fibrosis diseases although many of these genes were also identified targets in neurological disorders and ischemic stroke. Gene ontology analysis of the 167 common gene targets was largely unsurprising, though there was noted involvement in cellular metabolism, macromolecule biosynthesis, and cAMP, co-SMAD, and JNK-kinase activity (Supplemental Figure S6). MetaCore analysis of the 45 miRNA targets revealed four major pathways of interest which were regulated at several points by different identified miRNAs along the pathway (Figure 3E-H), suggesting deliberate targeting of certain pathways in a highly specific manner. The pathways of interest were the HNF4-a pathway (Figure 3E), with downstream regulation that promoted exosome trafficking and inhibited inflammation and apoptotic signaling^52–56^, the Smad3/STAT5 pathway (Figure 3F), with overall inhibition of signaling early in the Ang2 cardiac fibrosis cascade^57–62^, PPARg pathway/MAPK cascade (Figure 3G), with more multi-point regulation of Ang2-mediated fibrosis signaling and downstream targets parallel to the HNF4-a pathway^63–67^, and an SDF-1a/systemic stem cell recruitment pathway implicated in pro-regenerative healing from MI^68–73^ (Figure 4H).

**Figure 3.**
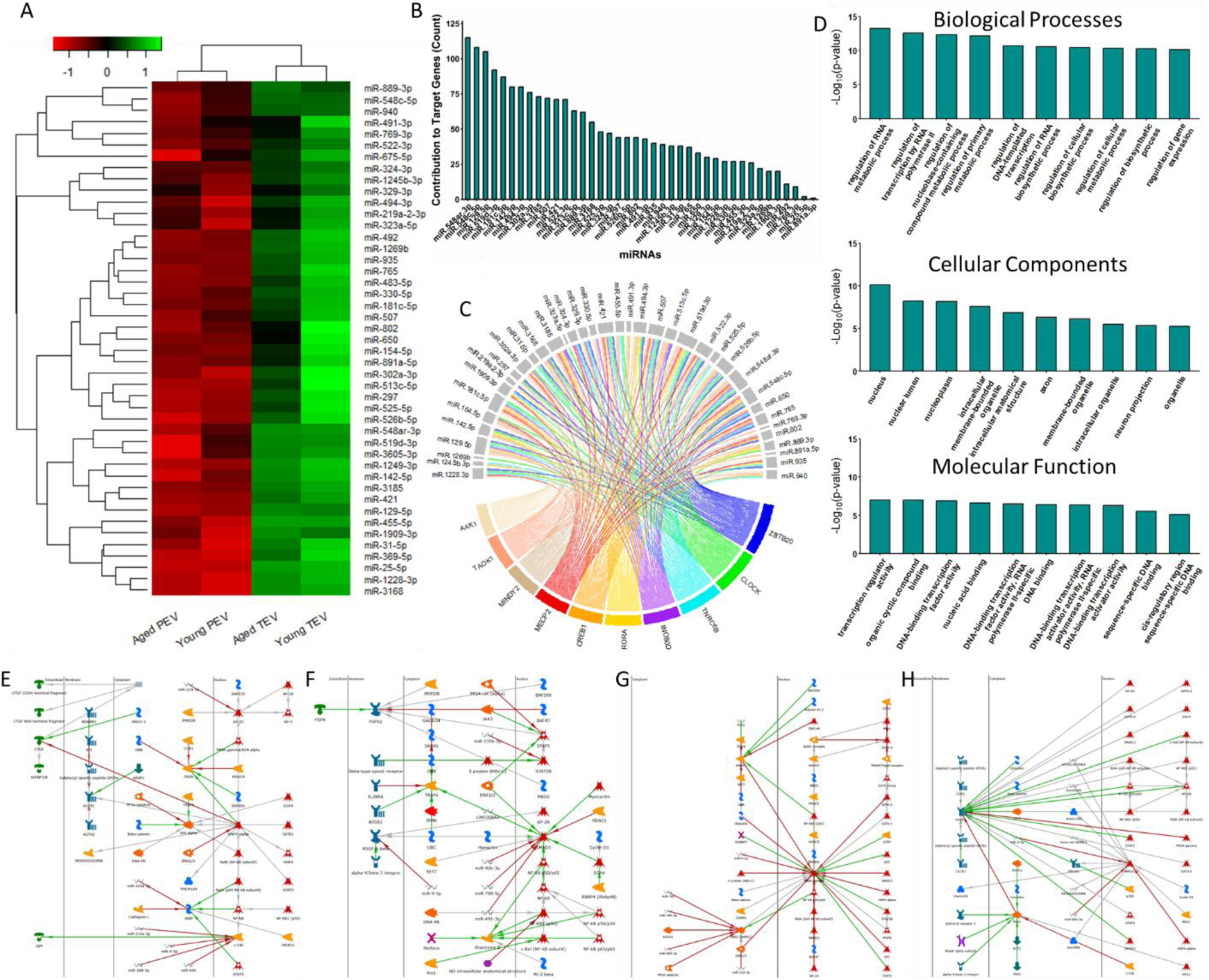
Cardiovascular-disease related target miRNAs and identified pathways identified from age and sex-related differences in sEV miRNA profile. Measurements were taken from n = 6 biological replicates per cohort, equally spread between male and female donors. (A) Heatmap showing identified target miRNAs enriched in TEVs relative to PEVs. (B) Analysis of miRNA contributors to predicted gene targets, showing by number the frequency of miRNAs contributing to different identified target genes (full gene list available in Supplemental Table S5). (C) Chord diagram showing overlap of miRNAs and the top 10 enriched predicted gene targets. (D) Gene ontology analysis of the predicted gene targets of the identified miRNAs showing the top 10 significant pathways identified (full graphs available in Supplemental Figure S6). (E-H) Results of MetaCore pathway analysis for the identified target miRNA data. * p < 0.05. Identified pathways were considered for analysis only for p-value < 0.05.

**Figure 4.**
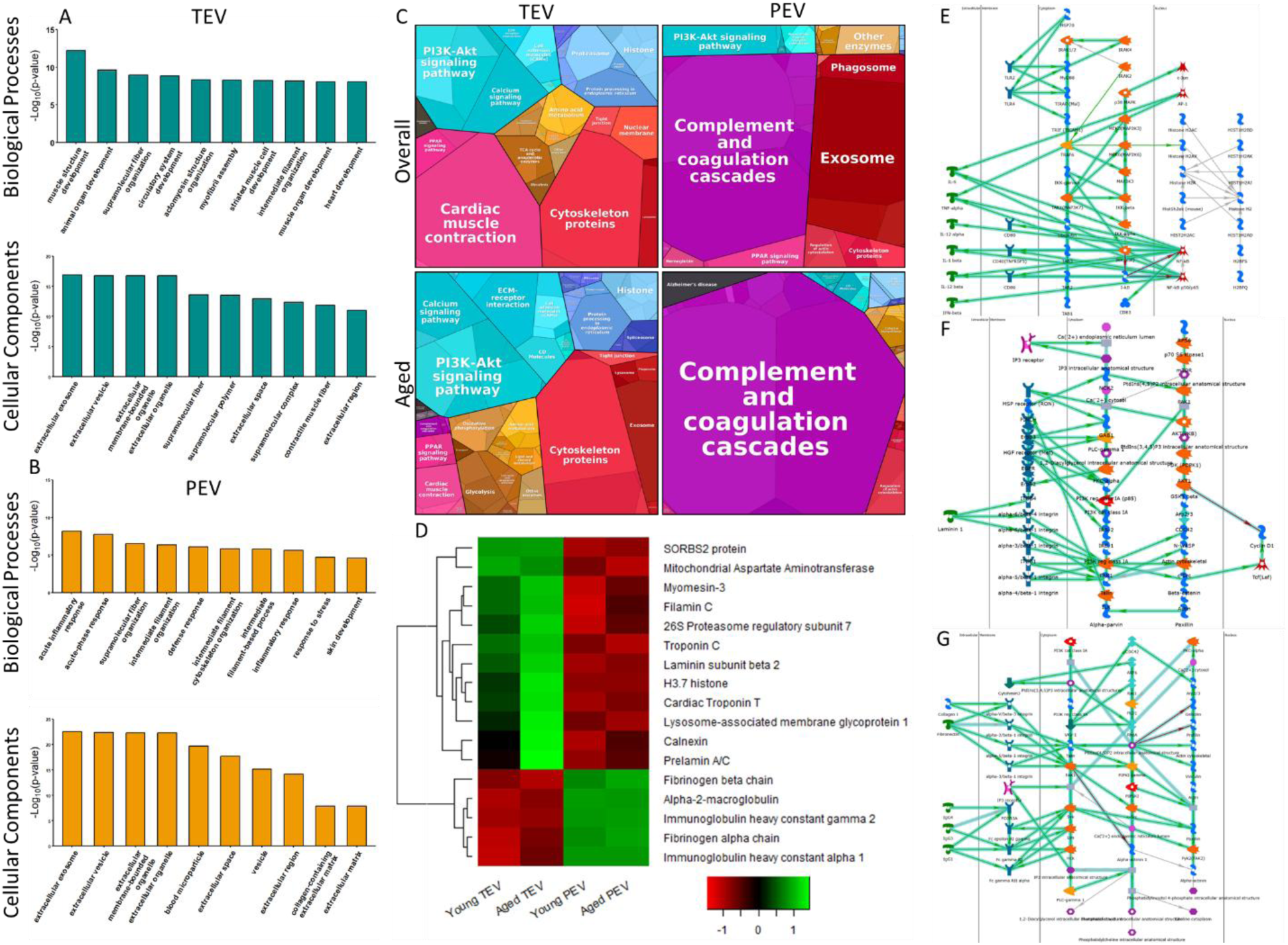
Cardiovascular-disease related target proteins and identified pathways identified from age-related differences in sEV protein contents. Measurements were taken from n = 6 biological replicates per cohort, equally spread between male and female donors. Gene Ontology Analysis results for TEVs (A) and PEVs (B) enriched in each population. Full graphs can be viewed at Supplemental Figure S9. (C) Proteomaps showing the KEGG pathways affected by the unique protein profiles of TEVs and PEVs overall, as well as for only aged donor TEVs and PEVs. Red: cellular processes; Yellow: metabolism; Pink: organismal systems; Blue: genetic information processing; Teal: environmental information processing; Purple: immune system; Black: human diseases. (D) Heatmap showing the 17 proteins of interest identified by supervised analysis of unique proteins for TEVs and PEVs. (E-G) Results of MetaCore pathway analysis for the unique protein profiles of TEVs (E, F) or PEVs (G). * p < 0.05. Identified pathways were considered for analysis only for p-value < 0.05.

### Gene ontology reveals functional differences in protein content between TEVs and PEVs

The proteins enriched in TEVs or PEVs were also assessed via gene ontology analysis (Figure 4A, B). Both TEV and PEV proteins were closely associated with extracellular vesicles and the extracellular space, as expected, however the associated biological processes predicted were notably different. TEVs were identified with muscular development and function (Figure 4A) as well as a number of pathways corresponding to specifically cardiac development and homeostasis and redox pathways (Supplemental Figure S9). PEV proteins, alternatively, corresponded more to acute inflammatory response and related pathways (Figure 4B), and was associated with IgG and other immunoglobulins and components (Supplemental Figure S9). Interestingly, the gene ontology analysis returned no significant results for molecular functions pertaining to PEV proteins.

### Tissue-resident sEVs showcase greater involvement in metabolism and homeostasis than plasma sEVs

Proteomap profiles (Figure 4C) were generated from the unique proteins identified for both TEVs (106 proteins) and PEVs (132 proteins) (Supplemental Table S8), as well as for aged donor TEVs (274 proteins) and PEVs (202 proteins) (Supplemental Table S9). Interestingly, overall TEV pathways were balanced between metabolism, cardiac muscle activity, and environmental signal transduction, whereas overall PEV pathways were dominated by immunoregulation and exosome handling. Overall TEVs also uniquely called a pathway for degenerative diseases (black, far left, top left proteomap). Aged donor sEV protein pathways, however, were fairly different. Aged donor TEVs showed less overall involvement in cardiac muscle activity and greater involvement in environmental signaling and metabolism. Aged donor TEVs, compared to overall, also became newly involved in immunoregulatory signaling as well as developing pathways associated specifically with cancer development, hypertrophic cardiomyopathy, and arrythmogenic right ventricular cardiomyopathy, as opposed to the previously observed degenerative diseases. Aged donor PEVs on the other hand became solely dominated by immunoregulatory signaling, excluding exosome handling almost entirely, and developing involvement in pathways associated with Alzheimer’s Disease. Potential individual protein targets were also identified from the unique proteins from TEVs and PEVs via supervised analysis (Figure 4D). These 17 proteins, while not necessarily directly implicated in major pathways, are common biomarkers for identification of disease state or other pathology and may allow for more rapid detection of critical marker quantification in clinical scenarios.

### Tissue-resident sEVs from older hearts are directly implicated in CVDs, but not plasma sEVs from older patients

The unique proteins from TEVs and PEVs were mapped via MetaCore pathway analysis, and the resulting pathways provided further insight into the downstream activities of TEVs (Figure 4E, F) and PEVs (Figure 4G). TEV proteins mediated NFkB/Interleukin signaling downstream of c-Jun and TRAF6 (Figure 4E), activities highly involved in cardiac immunoregulation and matrix homeostasis^74–78^, as well as directly interfacing with PI3k-1a, mTOR, and Laminin response (Figure 4F), which together can be implicated in response to cardiac fibrosis or other cardiac matrix remodeling^79–85^. Additionally, dysregulation of PI3k-1a and mTOR via the complex “PI3K/mTOR signaling pathway” network has been directly implicated in aging-related CVD onset and other aging-related pathologies, and the targeted inhibition of this pathway has reversed aging-related risk of post-MI cardiac fibrosis and pathological signaling post-cardiac insult^86,87^. PEVs also interacted with the surrounding matrix through integrin mediation (Figure 4G) but displayed more focus on modulating interactions with white blood cells via Igg manipulation, which may indicate a role in chronic inflammation in chronic CVD and heart failure^88–90^.

### Machine Learning Approaches Identify Novel Markers of Cardiac Aging

Variable selection techniques were applied to identify biomarkers that differentiate between young and aged cohorts for TEVs and PEVs and between TEV and PEV cohorts young and aged EVs (Figure 5A). This analysis revealed 42 potential miRNA targets for distinguishing between various cohorts (Supplemental Table S10). Of these, 5 miRNAs were targets which had previously been identified as potentially therapeutic miRNAs^7^, and one miRNA, miR-299, was a biomarker which could be collected from plasma which was indicative of cardiac aging.

**Figure 5.**
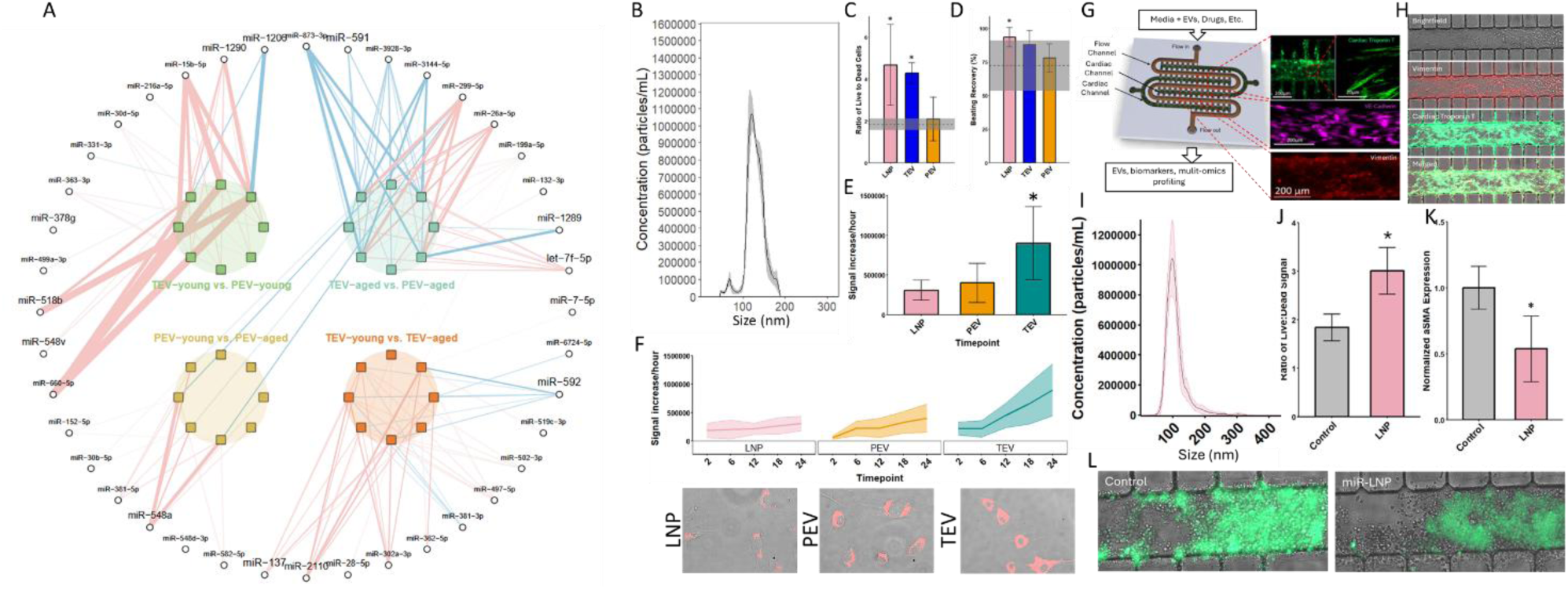
Machine-learning results inform novel theragnostic applications of EV-chaperoned miRNAs. For LNPs, n ≥ 3 in all assays; for TEVs and PEVs, n = 6 biological replicates per cohort, equally spread between male and female donors. A network plot representing the biomarkers selected from eight regularized logistic regression in 4 pairs of cohort comparisons: TEV-young vs. PEV-young; TEV-aged vs. PEV-aged; TEV-young vs. TEV-aged; PEV-young vs. PEV-aged, each depicted by a colored circle at the center of the network (A). The square nodes represent the eight models (with LASSO and SCAD penalties in one full dataset and three biologically defined subsets); the circular nodes along the periphery represent individual miRNA selected by at least one model. Edges connect miRNAs to the models in which they were selected, with red edges indicating a positive coefficient and blue edges indicating a negative coefficient. The thickness of each edge reflects the absolute magnitude of the coefficient. (B) Nanoparticle Tracking Analysis results for formulated lecithin lipid nanoparticles loaded with therapeutic miRNAs (miR-LNPs). Results for miR-LNP treatment of cells during 2 h hypoxia compared to TEVs and PEVs in CFs (C) and CMs (D). Uptake after 24 h (E) and uptake rate over time (F) for miR-LNPs compared to TEVs and PEVs. (G) Schematic of the <u>H</u>uman H<u>e</u>art <u>A</u>noxia and <u>R</u>eperfusion <u>T</u>issue (HEART) model with complex myocardial tissue and a perfusable vascular channel and (H) staining showing multi-cell type tissue in cardiac channel. (I) Nanoparticle tracking analysis for miR-LNPs administered to the HEART. LNP treatment resulted in (J) increased live:dead cell ratio and (K) decreased EMT after 2 h in MI-like conditions. (L) Representative stain of aSMA expression in the cardiac channel post-MI. * p < 0.05.

Interestingly, this analysis revealed only 21 potential protein targets for distinguishing between various cohorts (Supplemental Table S10). These were largely identified between young and aged PEVs, and aged TEVs and PEVs (Supplemental Figure S10).

### Treatment of Cardiac Cells with Therapeutic miRNAs Ameliorates Damage Under MI-like Conditions

Of the potentially therapeutic miRNA targets identified, three (miR-199a, miR-30d, miR-302a) were selected for testing with previously identified therapeutic miRNAs^7^. These miRNAs were encapsulated at 40 nM concentration in lecithin-based lipid nanoparticles (LNPs). The LNPs were ∼120 nm in size (Figure 5C), slightly smaller than both TEVs and PEVs, and looked as expected. Hypoxia assays on iCFs and iCMs compared to TEVs and PEVs revealed that miR-loaded LNPs (miR-LNPs) significantly increased iCF survival after hypoxia, comparable to TEVs (Figure 5D), with both showing more than 4-fold increase in living cells compared to the control. A similar, but less significant effect was observed in beating recovery, where TEVs showed a non-significant improvement but miR-LNPs showed a significant improvement compared to the control (Figure 5E). Uptake rate of the EVs was also assessed to determine if the effects were rate or concentration dependent, where TEVs demonstrated a significantly higher uptake rate than PEVs or miR-LNPs (Figure 5F). Furthermore, uptake rate increased over time for both PEVs and TEVs, but remained stagnant for miR-LNPs (Figure 5G).

## DISCUSSION

In this study, we isolated and characterized small extracellular vesicles (sEVs) isolated from patient blood plasma samples and from donor heart tissue samples. All sEV samples were taken from an equal number of men (n = 6 for tissue & plasma) and women (n = 6 for tissue and plasma) to minimize the impact of sex variance, and were bifurcated between aged (>50 y.o., n = 6 for tissue & plasma) and young (<40 y.o., n = 6 for tissue & plasma) for analysis (Supplemental Table S1). Both the plasma sEVs (PEVs) and tissue sEVs (TEVs) were observed under transmission electron microscopy (TEM) and population size distribution was assayed via nanoparticle tracking analysis (NTA), with mean and mode population sizes being consistent between both assays. Interestingly, the mode size of the sEV population differed with age in TEVs, but not PEVs, suggesting at least some degree of differential function with respect to aging between each category of EV. Surface protein composition was also assessed between aged and young PEVs and TEVs using western blot for classical exosome markers CD9, CD63, and TSG101^21^. While overall measured expression of these proteins increased with age in TEVs and decreased with age in PEVs, the relative expression of these proteins did not significantly change in either group, suggesting that the quantity, but not overall composition, of tetraspanin web targeting proteins changes with age, and that these changes are different in PEVs and TEVs. Finally, the miRNA and protein content of young and aged PEVs and TEVs were assayed. The miRNA content was measured using Nanostring technology, providing a full profile of human miRNAs, and unsupervised analysis was performed to identify distinguishing miRNAs between PEVs and TEVs which were independent of age. The TEV-specific age-independent miRNAs were further analyzed in MetaCore pathway analysis to provide a preliminary overview of processes uniquely modulated by TEVs. The protein content of all cohorts was similarly assessed using mass spectrometry, revealing 2547 proteins with varying expression levels across all biological replicates. Analysis of the dataset revealed 93 proteins consistently expressed in all biological replicates of all cohorts, with aged sEVs tending to have more commonly expressed proteins than young sEVs and TEVs tending to have more commonly expressed proteins than PEVs. Unique cohort proteins were also identified and assessed for downstream function using KEGG proteomapping software and MetaCore pathway analysis.

The size discrepancy in age-related size changes between TEVs and PEVs (Figure 1A, B) is an interesting result due to the known consistency of size distribution in EV populations^21^ and our previous study showing that these size differences are also not sex-dependent^7^. We first evaluated whether or not this discrepancy was due to different cell ages in the heart, as the difference due to cell age could have been mitigated in PEVs by the large number of organs which contribute to the PEV population. We evaluated this for both artificially aged iCFs (high passage cells at high confluency) and mCFs from aged mice (72 weeks), but found that neither population showed significantly different EV sizes produced compared to the young cells (Supplemental Figure S1). In previous studies, we have shown that the stiffness of the surrounding matrix can significantly alter the behavior of cells^10,13^, so an alternative cause may be differences in the matrix stiffness between young and aged hearts. This discrepancy will be further evaluated in future works to identify the precise relationship, if any, between matrix stiffness and produced EV size.

The uptake rate of EVs was also assessed for both populations and revealed novel interactions of TEVs and PEVs with tissue specificity, though only young PEVs and TEVs were used for this assay due to the substantially smaller size of aged TEVs compared to all other cohorts, which we have previously shown increases the uptake rate by cells^7^. To assess the specificity of TEVs and PEVs, uptake was evaluated for both populations with human and non-human (mouse) CFs. Interestingly, the uptake rate for both PEV groups and the non-human uptake of the TEVs was essentially the same at all timepoints measured (Figure 1F) but the uptake of TEVs on human CFs was significantly higher (Figure 1G) and in fact was increasing exponentially by the endpoint (Figure 1F). This suggests some level of specificity of the TEVs that is not present in the PEVs which may distinguish how EVs are selected to be exported to the plasma or to the surrounding ECM. The consistency of tetraspanin expression in TEV populations (Figure 1E) may be a partial contributor to this, as the tetraspanin web has been implicated in EV organ targeting systemically^39–41^, with several key, unidentified tetraspanins that are unique to TEVs contributing to the uniqueness of this specificity. Additionally, the exponential increase in rate of uptake of TEVs by human cells suggests that human cells taking up TEVs undergo some change which facilitates further TEV uptake. This has not yet been reported in literature and may be a key aspect of how TEVs provide therapeutic effects in young hearts, as an exponential increase in the uptake of therapeutic compounds over a short period of time would actively counteract increased inflammation and damage-associated effects common in the aftereffects of cardiac insult. These results demand further study, and will be more fully evaluated in future works specifically assessing the dynamics between uptake rate of natural or engineered EVs for the time-sensitive delivery of therapeutic agents.

The differential miRNA cargo of TEVs and PEVs provides an interesting avenue for the identification of therapeutic agents, as we have previously established that TEVs, but not PEVs, can promote anti-fibrotic effects in heart cells post-insult^7^. In light of this, the overlap in miRNA expression in the young PEVs and TEVs presents an interesting conundrum, although this is resolved by the nearly bifurcated miRNA expression in the aged EV populations (Figure 2A), suggesting that miRNA cargo substantially diverges between TEVs and PEVs with age. Despite this, it was possible to completely distinguish between TEVs and PEVs as a whole via 125 identified miRNAs selected by untargeted analysis of the entire miRNA profile for all cohorts (Figure 2C), providing further evidence that TEVs and PEVs are specialized for different functions. Of the 125 differentiating miRNAs, 59 were enriched in TEVs and 66 were enriched in PEVs, providing ample targets for further analysis of the specific function of each population. To better understand the unique cardiac-related function of the TEVs, we manually assessed the 59 TEV-enriched miRNAs for potential involvement in any CVD or CVD-related pathways via public databases and identified 45 targets for downstream analysis. MetaCore pathway analysis of the 45 revealed several pathways which were consistently regulated by several of the target miRNAs (Figure 3E-H). These combined effects served to inhibit targets in the HNF4-α pathway, Smad3/STAT5 pathway, and PPARγ pathway, and promote a therapeutic SDF-1α pathway. These results imply several exciting possibilities. First, we have previously hypothesized the cooperative effects of TEV miRNAs in promoting cardioprotective effects via synergistic inhibition of CVD-associated pathways^7^, but were only able to demonstrate this with 4 identified target miRNAs. The combined regulation of 3 damage associated pathways at multiple points by 45 target miRNAs provides a plethora of additional targets to evaluate in order to generate a TEV-mimetic cocktail of beneficial miRNAs for therapeutic applications. The use of a large number of miRNAs which endogenously cooperate to act on a small number of pathways would be able to provide much greater effect specificity than existing single-target drugs and previously proposed single-miRNA therapeutics. Additionally, these miRNAs together not only inhibit damage-associated effects but also promote reparative signaling. The SDF-1α is a known cardioprotective pathway which is difficult to activate without off-target effects. While the utility of these specific miRNAs requires additional study to determine efficacy and viability, this is an exciting first step in the development of highly specific therapeutics using endogenous agents.

The protein cargo of both EV populations also provided exciting results. While comparison of all identified proteins separated PEVs and TEVs, this result was not consistent across all biological replicates (Supplemental Figure S7). This was likely due to the inconsistency of protein expression across biological replicates (Figure 2D), with >60% of proteins being expressed in some, but not all, biological replicates of a given cohort. To be consistent in our analysis, we focused on proteins which were either expressed consistently in all cohorts (Figure 2E), or were uniquely expressed in a single cohort (Supplemental Table S8), although cohort-unique targets were still expressed in all biological replicates of a given cohort. Of the proteins consistently expressed in all cohorts, aged EVs tended to express more proteins overall for both TEVs and PEVs (Figure 2F), which is consistent with our previous hypothesis^7^. This suggests that overall proteins included in both systemic and tissue-resident increases with increasing age, which is consistent with the expected effects of inflammaging. Inflammaging has been identified as a potential cause of many aging-related pathologies, but is poorly defined and not well understood. The evaluation of EVs as part of aging-related increases in inflammation may provide a novel avenue for increased understanding of inflammaging and the development of intervention strategies.

Analysis of the proteins uniquely expressed in TEVs or PEVs via gene ontology and KEGG proteomapping revealed highly different functions of each population. PEVs were largely involved in immune system regulation (Figure 4B, 4C, Supplemental Figure S9), while TEVs were involved in a host of metabolic and regulatory functions in the heart (Figure 4A, 4C, Supplemental Figure S9). Interestingly, only TEVs were involved in disease progression (Figure 4C). To evaluate the aging-related changes in protein expression, the proteins unique to aged TEVs and PEVs were also mapped (Figure 4C). Again, PEVs were largely involved with the immune system, though immune system interactions made up a much larger proportion of involved pathways than PEVs overall, and now showed involvement with the onset of Alzheimer’s Disease pathways, which was an interesting pathway to remain consistent across all biological replicates. TEVs, however, showed much greater involvement in metabolism-related activities, less regulation of cardiac homeostasis, and new involvement in local inflammation and the direct onset of CVDs, differentiating aged TEVs from overall TEVs by direct implication in increased inflammation and disease progression. From the proteins uniquely expressed in TEVs or PEVs overall, 17 proteins of interest were identified which may be involved in the onset and progression of CVD. MetaCore analysis of these proteins revealed mulit-point involvement in three pathways which, while not directly implicated in CVD, are closely linked to the onset of several CVDs. These pathways included NFkB/Interleukin signaling downstream of c-Jun and TRAF6, PI3k-1a, mTOR, and Laminin response, which are each critical to regulating cardiac homeostasis and normal function. In particular, PI3k-1a and mTOR together are a regulatory pathway that is increasingly dysregulated with age, leading to increased risk of CVD, fibrosis, and heart failure^86,87^, while the dysregulation of Interleukin signaling and Laminin response pathways are closely related to the onset of chronic fibrosis and other pathological cardiac matrix remodeling ^79–85^. This data suggests that these pathways are, in some part, regulated by TEV signaling and that aging-related changes in TEV signaling, such as those we have shown, may disrupt these vital pathways and lead to increased risk of CVD. While more direct evaluation of the roles of TEVs on the regulation of these pathways is essential for understanding the complex relationship of aging-related EV changes and aging-related pathologies, this data provides a first step into understanding these processes.

Lastly, using the machine learning-based analyses we were able to validate several hypothesized therapeutic agents from our previous dataset^91^. In particular, miR-199a, which we had considered a significant agent in the initial therapeutic miRNA cocktail, was found to be enriched in young tissue, as we previously observed, and was identified by the analysis, in addition to miR-30d and miR-302a, which had been both identified as targets of interest (Figure 5A). To evaluate the therapeutic potential of these therapeutic miRNAs utilizing a drug delivery vehicle, the miRNAs were mixed at 40 nM and encapsulated in simple lecithin lipid nanoparticles (LNPs) (Figure 5B). While this miR-LNP treatment did significantly increase cell survival and recovery of function post-MI (Figure 5C, 5D), this function appeared to be rate-independent, as the uptake rate of LNPs was lower than both TEVs and PEVs yet the effect on cells was greater in both assays (Figure 5E, 5F). While the therapeutic cocktail will need additional formulation and testing, this is a promising early result for the development of a novel, endogenous anti-fibrotic therapy.

Overall, this study serves as an in-depth, comprehensive resource for identifying the cargo of sEVs isolated from plasma or cardiac tissue, as well as distinguishing between samples from young or aged individuals. These specific delineations are currently not well documented in literature, and increased assessment and validation of age-related and organ-specific paracrine signaling mechanisms will be essential in the development of next-generation therapeutics. Furthermore, these data suggest that, with increased understanding of both TEV and PEV interactions, it will be possible to closely investigate organ-specific health markers via plasma assays, one of the chief overall goals of biomarker research. The combination of novel machine learning and bioengineering approaches provides a unique avenue to achieve this goal in both a cost-efficient and broadly applicable manner for both other CVDs and other organs altogether. Finally, we demonstrate the feasibility of identifying novel, endogenous therapeutic agents by investigating the age-specific differences in TEV cargo, which we suggested in a previous publication^91^ The success of these novel therapeutics in ameliorating damage from MI-like conditions in cardiac cells opens the door to a new class of highly specific therapeutics which take advantage of endogenous repair machinery in the heart.

## MATERIALS & METHODS

### Human Heart Tissue Preparation

Donor human heart tissue was prepared according to our previous protocol^7^. Briefly, the left ventricle from frozen heart tissue was excised of fat and connective tissue and section into sub-300 µm slices, which were agitated in an aqueous acetic acid (0.1%, Sigma Aldrich, USA) and ethanol (4%, Sigma Aldrich, USA) solution at 200 rpm at room temperature (RT) for 2 hours. Tissues were then washed in sterile PBS for at least 30 minutes to remove free debris, then re-agitated in solution overnight at RT. Matrices were then vigorously washed in sterile PBS and sterile DI water, lyophilized, pulverized with pre-chilled mortar and pestle, and separated into 200 mg (dry weight) powder aliquots. ECM powder was resuspended in 1 mL of aqueous Collagenase Type II (0.2 mg/mL, Corning, USA), Tris buffer (50 mM, Sigma Aldrich, USA), and CaCl_2_ (5 mM, Amresco, USA) and mixed vigorously. Suspension was reacted overnight at RT with re-mixing every 6 h to prevent settling.

### Plasma Sample Collection

Whole blood was collected from patients via direct veinous puncture into ethylenediaminetetraacetic acid (EDTA)-treated tubes to prevent coagulation. Following collection, whole blood was centrifuged at 1000g and 4°C for 5 min to separate plasma. Patient plasma was then aliquoted to sterile RNAse-free tubes and shipped from the University of Florida College of Medicine to the University of Notre Dame at -80°C. After arrival, samples were stored at -80°C until use.

### Vesicle Extraction and Isolation

For both digested heart tissue ECM and collected plasma samples, the solution was centrifuged three times at 500g for 10 minutes, 2500g for 20 minutes, and 10,000g for 30 minutes. The pellet was discarded after each centrifugation to remove any insoluble contaminants. After the final centrifugation, the collected supernatant was ultracentrifuged at 100,000g for 70 minutes at 4°C using an ultracentrifuge (Optima MAX-XP Tabletop Ultracentrifuge, Beckman Coulter). The resulting pellet from ultracentrifugation was either used immediately or stored dry at -80°C.

### Transmission Electron Microscopy

Single pellets were fixed via resuspension in 2.5% glutaraldehyde solution for 2 h at RT in the dark. Fixed samples were then loaded onto plasma-cleaned Formvar/carbon-coated copper 200 mesh grids (Polysciences) and negative-stained with Vanadium staining solution (Abcam, ab172780). The resulting grids were imaged at 80 kV with a TEM (JEOL 2011, Japan).

### Nanoparticle Tracking Analysis

Single pellets were resuspended in 500 µL of sterile, particle-free PBS and measured using a NanoSight NS300 machine (Malvern Panalytical) with NTA software version 3.2.16. Resuspended samples were kept at 4 °C until measurement, and measurements were taken at RT.

### Western Blot

Single pellets were lysed in RIPA buffer containing 1% proteinase inhibitor cocktail (Brand, Country) at 4°C for 30 minutes and resultant protein concentration was measured via bicinchoninic acid (BCA) assay (Pierce Chemical). Equal amounts of protein for each sample were separated by 12% SDS-PAGE and transferred to blotting membranes. Blotting membranes were incubated on a rocker overnight at 4°C with rabbit polyclonal primary antibodies against CD9 (Abcam, ab223052), CD63 (Abcam, ab216130), and TSG101 (Abcam, ab30871) at (1:1000), then for 1 h at RT with HRP-conjugated goat anti-rabbit secondary antibody (Abcam, ab205718). Stained membranes were then treated with a chemiluminescent substrate (Clarity ECL, Bio-Rad) and imaged using a ChemiDoc-It2 imager (UVP, Analytik Jena) equipped with VisionWorks software. Images were processed using ImageJ (NIH).

### Culture of Human Induced Pluripotent Stem Cells (hiPSCs)

DiPS 1016 SevA hiPSCs, which were derived from human skin fibroblasts, were cultured on Geltrex (1% Invitrogen, USA)-coated culture flasks in mTeSR (StemCell Technologies, Canada) media supplemented with 1% penicillin (VWR, USA) under standard culture conditions from passages 40-50.

Cells were passaged at 80% confluency. To passage the hiPSCs, cells were detached using Accutase (StemCell Technologies, Canada) and seeded onto Geltrex-coated cell culture well plates or split between cell culture flasks. Seeding was performed in mTeSR media supplemented with Rho-associated, coiled-coil containing protein kinase (ROCK) inhibitor (5 μM, StemCell Technologies, Canada). For differentiation, cells were cultured until 95% confluency before starting a protocol

### Differentiation and Culture of hiPSC-derived Cardiac Fibroblasts (iCFs)

In-house differentiated induced pluripotent stem cell (iPSC)-derived cardiac fibroblasts (iCFs) were generated and cultured according to our previous protocol^13^, and were seeded on fibronectin (50 µL/mL) -coated 24-well cell culture plates. Mouse primary cardiac fibroblasts were isolated from collected mouse hearts according to our previous protocol^7^, and were seeded on fibronectin-coated 24-well cell culture plates as with iCFs. Both iCFs and mCFs were cultured in Dulbecco’s Modified Eagle’s Medium (DMEM, Thermo Fisher, USA), supplemented with fetal bovine serum (FBS, 10%, Gibco, USA), penicillin/streptomycin antibiotics (P/S, 1%, Life Technologies, USA), and SD208 (3 µM, Sigma Aldrich, USA).

### Culture of Mouse Primary Cardiac Fibroblasts (mCFs)

Isolated mCFs were seeded on fibronectin-coated 24-well cell culture plates as with iCFs. Both iCFs and mCFs were cultured in Dulbecco’s Modified Eagle’s Medium (DMEM, Thermo Fisher, USA), supplemented with fetal bovine serum (FBS, 10%, Gibco, USA), penicillin/streptomycin antibiotics (P/S, 1%, Life Technologies, USA), and SD208 (3 µM, Sigma Aldrich, USA).

### Cellular Uptake of sEVs

Cellular uptake of sEVs was performed based on our previous protocol^7^. Briefly, sEV pellets were resuspending in PBS and concentration was assessed via BCA assay and subsequently stained using the ExoGlow-Membrane staining kit (System Biosciences, USA) for 30 min at RT. Stained sEVs or a blank control were then purified in a salt gradient column for a final sEV concentration of 25 µg/mL in DMEM, supplemented with exosome-free FBS (10%) and 1% P/S. Cells were incubated with the sEV-conditioned media at 37°C and imaged at 0, 3, 8, 16, and 24 h incubation. During imaging the conditioned media was exchanged with sterile, particle-free PBS, and the same media was re-placed in the same wells immediately preceding return to incubation.

### miRNA Isolation

Internal RNAs were isolated from sEVs using the Total Exosome RNA & Protein Isolation Kit (Thermo Fisher Scientific, USA) according to the manufacturer’s protocol. Briefly, single pellets were resuspended in sterile, particle-free PBS and incubated with an equal volume of provided denaturation buffer 4 °C for 5 min. The solution was then mixed with an equal volume of Acid-Phenol:Chloroform by vortexing for 30 seconds and centrifuged for 5 min at 15,000g. The resulting aqueous phase was extracted and combined with 1.25x volume of 100% ethanol, then transferred to the provided spin column. The spin column was centrifuged at 10,000g for 15 seconds to bind and wash the RNA, then the RNA was eluted in 100 µL of the provided elution solution. Eluted RNA was then concentrated using 3 kDa microcentrifuge spin filters (Amicon, USA). 100 µL miRNA solution was worked up to 420 µL with RNAse-free water and placed into a filter, then centrifuged at 14,000g for 90 minutes. Next, the filter was inverted into a fresh collection tube, and centrifuged at 8,000g for 2 minutes. The resulting 20-25 µL of RNA concentrate was immediately quantified by microvolume spectrophotometery (Nanodrop 2000, Thermo Fisher Scientific, USA).

### Profiling of Total miRNA Population

Concentrated RNA isolate was prepared for Nanostring miRNA profiling according to the manufacturer’s protocol. Briefly, the provided miRNA codeset was mixed with the provided hybridization buffer to produce a master mix, and spike-in miRNA controls were prepared at 200 pm. In order, the master mix, concentrated sample miRNA, spike-in miRNA, and provided probes were mixed in a PCR plate and incubated at 65 °C for 16 h. The hybridized solution was then mixed with 15 µL of provided hybridization buffer, for a total volume of 30-35 µL, and added to the provided microfluidic cartridge. The assay was run with the provided protocol for total miRNA analysis, and data was processed and analyzed using the provided software using the recommended settings. The dataset was then exported for further analysis and target selection.

### miRNA Gene Target Identification and Gene Ontology

Likely gene targets of miRNAs were predicted by miRDB^92^, miRTarBase^93^, and TargetScan^94^, and commonly predicted genes were compiled for further analysis (Supplemental Table S5). In-house software was used to identify unique genes and match miRNAs to the corresponding gene targets to determine overlapping gene targets between miRNAs of interest. Gene targets were selected as genes which were predicted targets for greater than or equal to 10 miRNA targets (Supplemental Table S6).

Gene Ontology Analysis (GOA) was performed on the identified gene targets using an online database via PANTHER Gene Ontology classification for enrichment analysis^95–97^. Data was extracted from the output and graphed using R.

### miRNA Pathway Analysis

Exported Nanostring data was normalized via Log10 normalization and uploaded to the Clarivate MetaCore system for pathway analysis, then cleaned to remove erroneous miRNA detection, if any, by the system. Automated network analysis was conducted with 50 nodes per network. Results were presented as pathways obtained from the software.

### Protein Isolation

Single pellets were lysed in 1% Triton-X solution at 4°C overnight, and resultant protein concentration was measured via bicinchoninic acid (BCA) assay (Pierce Chemical, USA). The resulting protein solution was solubilized in an SDS buffer solution consisting of 5% SDS, 50 nM TEAB (pH 7.55) at room temperature until fully combined.

### Total Protein Profiling

Total protein profiling was performed via mass spectrometery (Q Exactive HF Mass Spectrometer) according to the manufacturer’s protocol. Briefly, solubilized protein samples were denatured in 20 mM DTT, then alkylated with 40 mM iodoacetamide. Alkylated samples were digested in a provided column with trypsin (1:25 wt:wt) in 50 mM TEAB (pH 8) at 37°C overnight. Digested samples were then desalted via ziptip C18 and dried, then reconstituted in acetonitrile. Mass spectrometry was run with 1 µL of acetonitrile-reconstituted solution for bottom-up proteomics.

### Protein pathway mapping

GOA was performed on enriched proteins using the same protocol as described above. Data was extracted from the output dataset and graphed using R. Proteomic interactions of uniquely enriched proteins were classified through KEGG-based proteomapping software^98^ and are presented as obtained.

### Machine Learning-Based Analysis

The datasets analyzed consisted of four cohorts: TEV-young (n=6), TEV-aged (n=6), PEV-young (n=6), and PEV-aged (n=6) for the analysis of 798 identified miRNAs; TEV-young (n=6), TEV-aged (n=6), PEV-young (n=6), and PEV-aged (n=6) for the analysis of 2547 identified proteins. Regularized logistic regression was used to identify biomarkers potentially associated with cohort differences. Comparisons were performed between young (TEV-young vs. PEV-young), aged (TEV-aged vs. PEV-aged), TEV (TEV-young vs. TEV-aged), and PEV (PEV-young vs. PEV-aged) groups.

The negative log-likelihood function for the logistic regression is:

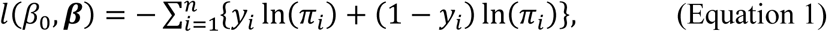

where the log-odds are

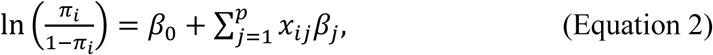

for *i* = 1, 2, …, *n*, and ***x****_i_* is a *p* × 1 vector that contains the values of *p* miRNA biomarkers in sample *i,* and *y_i_* ∈ {0, 1} is a binary group indicator. *β*_0_ is the intercept and ***β*** = (*β*_1_, *β*_2_, … *β_p_*) is a *p* × 1 vector of regression coefficients. To perform variable selection, a penalty term was added to the likelihood:

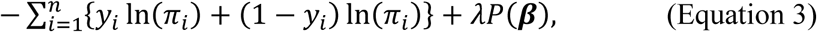

where the penalty term *λP*(***β***) represents the product of tuning parameter *λ* and the penalty function *P*(***β***). Penalized regression is often used for variable selection and regularizing parameter estimation and predictions when *p* is large, especially when *n* is small, so as to improve the robustness and generalizability of the learned model.

Two widely adopted penalty functions were applied: LASSO (least absolute shrinkage and selector operation)^99^ and SCAD (smoothly clipped absolute deviation)^100^. The penalty functions are defined as:

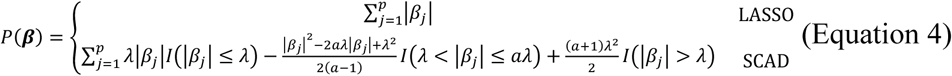

The tuning parameter 𝜆 was selected via the leave-one-out cross validation (LOOCV).

In addition to running the above logistic regression models with LASSO and SCAD penalties on the full datasets (798 miRNAs, 2547 proteins), the analyses were also performed on biologically informed subsets. The miRNA dataset was divided into 3 subsets: (1) purely cardiac specific miRNAs (450); (2) cardiac specific and vasculature miRNAs (565); (3) cardiac specific, vasculature, cerebral vasculature, and rare cardiac disease miRNAs (576) – leading to 8 models in total for each comparison. For the protein dataset, subsets were created based on proteins with non-zero values across all biological replicates in a given comparison (young: 127, aged: 220, tissue: 209, plasma: 225). Targets were selected from biomarkers identified in the subsetted analysis.

All analyses were performed in R. For LASSO regression, R package “glmnet v4.1-8” was used; for SCAD, R package “ncvreg v3.14.3” was employed.

Regarding the coefficient interpretation, each logistic regression model uses a binary variable to indicate group membership, where the reference group is coded as 0 (e.g., **aged** or **TEV**), and the comparison group is coded as 1 (e.g., **young** or **PEV**). The model estimates the log-odds of belonging to the comparison group based on the expression levels of selected biomarkers. A **positive coefficient** implies that increased expression of the biomarker is associated with higher odds of being in the comparison group (e.g., young or PEV), whereas a **negative coefficient** indicates that the biomarker is more strongly associated with the reference group (e.g., aged or TEV).

### Lipid Nanoparticle (LNP) Formation and miRNA Encapsulation

Lipid nanoparticles (LNPs) formed from lecithin (soy, Neogen), an inexpensive and benign lipid^101^ of recent interest in synthesis of LNPs from naturally occurring lipids^102^, particularly for cardiac drug delivery^103^, were formed by simple hot extrusion. Briefly, mixed into PBS at 3 mM concentration and heated to 50℃ while mixing to form a lipid emulsion. This temperature is sufficient for lecithin mixing into water for LNP formation^104^ and low enough to avoid denaturing miRNAs during lipid nanoparticle (LNP) formation^105^. Once lecithin is sufficiently mixed, miRNA mimics were added to the solution for final concentration of 40 nM total miRNAs. The miRNA mimics added were miR-199a (ACAGUAGUCUGCACAUUGGUUA, miRVana), miR-30d (UGUAAACAUCCCCGACUGGAAG, miRVana), miR-302a (ACUUAAACGUGGAUGUACUUGCU, miRVana) miR-125 (ACGGGUUAGGCUCUUGGGAGCU, miRVana), miR-143 (UGAGAUGAAGCACUGUAGCUC, miRVana), and miR-222 (AGCUACAUCUGGCUACUGGGU, miRVana). Once the miRNAs were added, the lipid-miR mixture was hot extruded^106^ while cooling, through a 100 nm membrane to generate LNPs with miRNAs encapsulated (miR-LNPs). This was done using a Mini-Extruder (Avanti Polar Lipids) and corresponding membrane.

### Hypoxia Assay

To analyze the contractility of iCMs, a block-matching algorithm was performed using MATLAB as described previously^107^. Briefly, iCM or 3D structures were recorded in brightfield in real time under a microscope (Axio Observer. Z1, Zeiss, Hamamatsu C11440 digital camera) for 30 s intervals. Videos were then uploaded to the analysis software, and beating velocity and frequency were calculated, and contraction heat maps were plotted. Baseline beating rates of iCMs were measured before beginning the hypoxia assay.

iCFs or iCMs were transferred to deoxygenated (>12 h hypoxia) media containing 1 milliong particle/mL LNPs with or without miRNA encapsulation (miR-LNP and control, respectively), or TEV or PEVs at 25 µg/mL as measured by bicinchoninic acid (BCA) assay (Pierce Chemical). Cell were then subjected to hypoxia for 3 hours, which we have previously shown is sufficient to induce MI-like cell death^6^. Following hypoxia, iCF plates were stained with calcein AM and ethidium homodimer-1 Live/Dead stains (Invitrogen, 1:1000) to quantify living and dead cells, respectively. Stained cells were imaged, and images were quantified in ImageJ.

To analyze the contractility of iCMs, a block-matching algorithm was performed using MATLAB as described previously^107^. Briefly, iCM or 3D structures were recorded in brightfield in real time under a microscope (Axio Observer. Z1, Zeiss, Hamamatsu C11440 digital camera) for 30 s intervals. Videos were then uploaded to the analysis software, and beating velocity and frequency were calculated, and contraction heat maps were plotted, and % recovery was calculated relative to the control. Timepoints were taken every 12 h following the removal of the iCMs from hypoxia.

### Statistical Analysis

Results were analyzed by one-way analysis of variance (ANOVA) with post-hoc Tukey’s HSD, two-way ANOVA with post-hoc Tukey’s multiple comparison test, or a two-tailed Welch t-test for unequal standard deviation. Values are presented as the mean ± standard deviation (SD) unless otherwise indicated, and differences were considered significant when p ≤ 0.05.

## List of Supplementary Materials

Fig S1 to S10

Table S1 to S10

## ACKNOWLEDGEMENTS

We thank Dr. Kieth March for providing the plasma samples utilized in this study.

The lyophilization of decellularized ECM was conducted at the Center for Environmental Science and Technology (CEST) at the University of Notre Dame.

We thank the Biophysics Instrumentation (BIC) Core Facility for the use of Optima MAX-XP Tabletop Ultracentrifuge.

The authors acknowledge the use of the Electron Microscopy Core of the Notre Dame Integrated Imaging Facility, a designated core of the NIH-funded Indiana Clinical and Translational Sciences Institute.

The Nanoparticle Tracking Analysis was conducted using the NanoSight NS300 at the Harper Cancer Research Institute (HCRI) Tissue Core Facility.

## Data Availability Statement

All data required for production of the manuscript is included in this submission. Additional raw data can be provided upon request.

## Ethics Statement

Human heart tissue was collected from donors whose hearts were deemed unsuitable for transplantation through the Indiana Donor Network (IDN) or National Disease Research Institute (NDRI). IRB approval was waived, as no identifying information was provided by the Indiana Donor Network or NDRI. De-identified human whole blood was collected from live donors at the University of Florida College of Medicine through an IRB approved standard collection protocol (IRB#201901232). All tissue collection was performed in accordance with the declaration of Helsinki.

## REFERENCES

1. Rahimi, K., Duncan, M., Pitcher, A., Emdin, C. A. & Goldacre, M. J. Mortality from heart failure, acute myocardial infarction and other ischaemic heart disease in England and Oxford: a trend study of multiple-cause-coded death certification. J Epidemiol Community Health 69, 1000–1005 (2015).

2. Reyes-Farias, M., Fos-Domenech, J., Serra, D., Herrero, L. & Sánchez-Infantes, D. White adipose tissue dysfunction in obesity and aging. Biochem Pharmacol 192, 114723 (2021).

3. Laconi, E., Marongiu, F. & DeGregori, J. Cancer as a disease of old age: changing mutational and microenvironmental landscapes. British Journal of Cancer 2020 122:7 122, 943–952 (2020).

4. Xi, J. Y., Lin, X. & Hao, Y. T. Measurement and projection of the burden of disease attributable to population aging in 188 countries, 1990-2050: A population-based study. J Glob Health 12, (2022).

5 Population Reference Bureau (PRB). Countries With the Oldest Populations in the World. https://www.prb.org/resources/countries-with-the-oldest-populations-in-the-world/ (2020).

6. Ozcebe, S. G., Bahcecioglu, G., Yue, X. S. & Zorlutuna, P. Effect of cellular and ECM aging on human iPSC-derived cardiomyocyte performance, maturity and senescence. Biomaterials 268, 120554 (2021).

7. Ronan, G., Bahcecioglu, G., Yang, J. & Zorlutuna, P. Age and Sex-Dependent Differences in Human Cardiac Matrix-Bound Exosomes Modulate Fibrosis through Synergistic miRNA Effects. bioRxiv 2022.11.14.516464 (2022) doi:10.1101/2022.11.14.516464.

8. Ozcebe, S. G. & Zorlutuna, P. In need of age-appropriate cardiac models: Impact of cell age on extracellular matrix therapy outcomes. Aging Cell 22, e13966 (2023).

9. Basara, G. et al. Myocardial infarction from a tissue engineering and regenerative medicine point of view: A comprehensive review on models and treatments. Biophys Rev 3, 031305 (2022).

10. Basara, G., Gulberk Ozcebe, S., Ellis, B. W. & Zorlutuna, P. Tunable Human Myocardium Derived Decellularized Extracellular Matrix for 3D Bioprinting and Cardiac Tissue Engineering. Gels 2021, Vol. 7, Page 70 **7**, 70 (2021).

11. Yang, J., Bahcecioglu, G. & Zorlutuna, P. The Extracellular Matrix and Vesicles Modulate the Breast Tumor Microenvironment. Bioengineering (Basel) 7, (2020).

12. Ellis, B. W. et al. Adipose Stem Cell Secretome Markedly Improves Rodent Heart and hiPSC-derived Cardiomyocyte Recovery from Cardioplegic Transport Solution Exposure. Stem Cells 39, 170 (2021).

13. Basara, G., Celebi, L. E., Ronan, G., Discua Santos, V. & Zorlutuna, P. 3D bioprinted aged human post-infarct myocardium tissue model. Health Sci Rep 7, e1945 (2024).

14. Ozcebe, S. G., Tristan, M. & Zorlutuna, P. Adult Human Heart ECM Improves Human iPSC-CM Function via Mitochondrial and Metabolic Maturation. bioRxiv 2023.10.31.565062 (2023) doi:10.1101/2023.10.31.565062.

15. Ellis, B. W. et al. Human Heart Anoxia and Reperfusion Tissue (HEART) Model for the Rapid Study of Exosome Bound miRNA Expression As Biomarkers for Myocardial Infarction. Small 18, 2201330 (2022).

16. Ellis, B. W., Acun, A., Isik Can, U. & Zorlutuna, P. Human iPSC-derived myocardium-on-chip with capillary-like flow for personalized medicine. Biomicrofluidics 11, 024105 (2017).

17. Ronan, G., Bahcecioglu, G., Aliyev, N. & Zorlutuna, P. Engineering the cardiac tissue microenvironment. Progress in Biomedical Engineering 6, 012002 (2023).

18. D’Anca, M. et al. Exosome Determinants of Physiological Aging and Age-Related Neurodegenerative Diseases. Front Aging Neurosci 11, 232 (2019).

19. Eitan, E. et al. Age-Related Changes in Plasma Extracellular Vesicle Characteristics and Internalization by Leukocytes. Sci Rep 7, 1342 (2017).

20. Sheta, M., Taha, E. A., Lu, Y. & Eguchi, T. Extracellular Vesicles: New Classification and Tumor Immunosuppression. Biology (Basel) 12, (2023).

21. Welsh, J. A. et al. Minimal information for studies of extracellular vesicles (MISEV2023): From basic to advanced approaches. J Extracell Vesicles 13, e12404 (2024).

22. Xu, M. Y., Ye, Z. S., Song, X. T. & Huang, R. C. Differences in the cargos and functions of exosomes derived from six cardiac cell types: a systematic review. Stem Cell Res Ther 10, 194 (2019).

23. Yáñez-Mó, M. et al. Biological properties of extracellular vesicles and their physiological functions. J Extracell Vesicles 4, 27066 (2015).

24. Jia, Y. et al. Small extracellular vesicles isolation and separation: Current techniques, pending questions and clinical applications. Theranostics 12, 6548 (2022).

25. Huang, K. et al. An off-the-shelf artificial cardiac patch improves cardiac repair after myocardial infarction in rats and pigs. Sci Transl Med 12, (2020).

26. Ren, X. et al. A multiplexed ion-exchange membrane-based miRNA (MIX·miR) detection platform for rapid diagnosis of myocardial infarction. Lab Chip 21, 3876–3887 (2021).

27. Bahcecioglu, G. et al. Aged Breast Extracellular Matrix Drives Mammary Epithelial Cells to an Invasive and Cancer-Like Phenotype. bioRxiv 2020.09.30.320960 (2020) doi:10.1101/2020.09.30.320960.

28. Yang, J., Bahcecioglu, G., Ronan, G. & Zorlutuna, P. Aged breast matrix bound vesicles promote breast cancer invasiveness. Biomaterials 306, 122493 (2024).

29. Basara, G. et al. Electrically conductive 3D printed Ti3C2Tx MXene-PEG composite constructs for cardiac tissue engineering. Acta Biomater 139, 179 (2022).

30. Law, M. R., Watt, H. C. & Wald, N. J. The underlying risk of death after myocardial infarction in the absence of treatment. Arch Intern Med 162, 2405–2410 (2002).

31. Cupples, L. A., Gagnon, D. R., Wong, N. D., Ostfeld, A. M. & Kannel, W. B. Preexisting cardiovascular conditions and long-term prognosis after initial myocardial infarction: The Framingham Study. Am Heart J 125, 863–872 (1993).

32. Narins, C. R. et al. Relationship Between Intermittent Claudication, Inflammation, Thrombosis, and Recurrent Cardiac Events Among Survivors of Myocardial Infarction. Arch Intern Med 164, 440–446 (2004).

33. Zhu, J. et al. The incidence of acute myocardial infarction in relation to overweight and obesity: a meta-analysis. Arch Med Sci 10, 855 (2014).

34. Yusuf, S. et al. Effect of potentially modifiable risk factors associated with myocardial infarction in 52 countries (the INTERHEART study): case-control study. Lancet 364, 937–952 (2004).

35. Florio, M. C., Magenta, A., Beji, S., Lakatta, E. G. & Capogrossi, M. C. Aging, MicroRNAs, and Heart Failure. Curr Probl Cardiol 45, 100406 (2020).

36. Anand, S. S. et al. Risk factors for myocardial infarction in women and men: insights from the INTERHEART study. Eur Heart J 29, 932–940 (2008).

37. Piccinini, A. M. & Midwood, K. S. Illustrating the interplay between the extracellular matrix and microRNAs. Int J Exp Pathol 95, 158–180 (2014).

38. Huleihel, L. et al. Matrix-bound nanovesicles within ECM bioscaffolds. Sci Adv 2, (2016).

39. Smith, Z. J. et al. Single exosome study reveals subpopulations distributed among cell lines with variability related to membrane content. J Extracell Vesicles 4, 28533 (2015).

40. Rana, S., Yue, S., Stadel, D. & Zöller, M. Toward tailored exosomes: The exosomal tetraspanin web contributes to target cell selection. Int J Biochem Cell Biol 44, 1574–1584 (2012).

41. Barranco, I. et al. Extracellular vesicles isolated from porcine seminal plasma exhibit different tetraspanin expression profiles. Scientific Reports 2019 9:1 9, 1–9 (2019).

42. Li, F. et al. Zinc Finger Protein ZBTB20 protects against cardiac remodelling post-myocardial infarction via ROS-TNFα/ASK1/JNK pathway regulation. J Cell Mol Med 24, 13383–13396 (2020).

43. Corella, D. et al. CLOCK gene variation is associated with incidence of type-2 diabetes and cardiovascular diseases in type-2 diabetic subjects: Dietary modulation in the PREDIMED randomized trial. Cardiovasc Diabetol 15, 1–12 (2016).

44. Edwards, T. L., Michels, K. A., Hartmann, K. E. & Velez Edwards, D. R. BET1L and TNRC6B associate with uterine fibroid risk among European Americans. Hum Genet 132, 943–953 (2013).

45. Bai, Z., Luo, Y. & Tian, L. ERCC5, HES6 and RORA are potential diagnostic markers of coronary artery disease. FEBS Open Bio 12, 1814–1827 (2022).

46. Zhang, T. & Ge, J. Mechanism of CREB1 in cardiac function of rats with heart failure via regulating the microRNA-376a-3p/TRAF6 axis. Mammalian Genome 33, 490–501 (2022).

47. Wang, C. et al. The Effect of Mecp2 on Heart Failure. Cellular Physiology and Biochemistry 47, 2380–2387 (2018).

48. Bone, W. P. et al. Multi-trait association studies discover pleiotropic loci between Alzheimer’s disease and cardiometabolic traits. Alzheimers Res Ther 13, 1–14 (2021).

49. Fang, C. Y., Lai, T. C., Hsiao, M. & Chang, Y. C. The Diverse Roles of TAO Kinases in Health and Diseases. International Journal of Molecular Sciences 2020, Vol. 21, *Page* 7463 21, 7463 (2020).

50. Song, Y. et al. M2 Microglia Extracellular Vesicle miR-124 Regulates Neural Stem Cell Differentiation in Ischemic Stroke via AAK1/NOTCH. Stroke 54, 2629–2639 (2023).

51. Xin, X. et al. Development and therapeutic potential of adaptor-associated kinase 1 inhibitors in human multifaceted diseases. Eur J Med Chem 248, 115102 (2023).

52. Rafiq, K. et al. C-Cbl inhibition improves cardiac function and survival in response to myocardial ischemia. Circulation 129, 2031–2043 (2014).

53. Tang, Z. et al. Suppression of c-Cbl tyrosine phosphorylation inhibits neointimal formation in balloon-injured rat arteries. Circulation 118, 764–772 (2008).

54. De Melker, A. A., Van Der Horst, G., Calafat, J., Jansen, H. & Borst, J. c-Cbl ubiquitinates the EGF receptor at the plasma membrane and remains receptor associated throughout the endocytic route. J Cell Sci 114, 2167–2178 (2001).

55. Cormont, M. et al. CD2AP/CMS Regulates Endosome Morphology and Traffic to the Degradative Pathway Through its Interaction with Rab4 and c-Cbl. Traffic 4, 97–112 (2003).

56. Ravandi, A. et al. Relationship of IgG and IgM autoantibodies and immune complexes to oxidized LDL with markers of oxidation and inflammation and cardiovascular events: Results from the EPIC-norfolk study. J Lipid Res 52, 1829–1836 (2011).

57. Zheng, Y. et al. Modulation of STAT3 and STAT5 activity rectifies the imbalance of Th17 and Treg cells in patients with acute coronary syndrome. Clinical Immunology 157, 65–77 (2015).

58. Kurdi, M., Zgheib, C. & Booz, G. W. Recent Developments on the Crosstalk Between STAT3 and Inflammation in Heart Function and Disease. Front Immunol 9, 421289 (2018).

59. Pang, Q. et al. Regulation of the JAK/STAT signaling pathway: The promising targets for cardiovascular disease. Biochem Pharmacol 213, 115587 (2023).

60. Liu, J. ; et al. The Role of JAK/STAT Pathway in Fibrotic Diseases: Molecular and Cellular Mechanisms. Biomolecules 2023, Vol. 13, Page 119 13, 119 (2023).

61. Hanna, A., Humeres, C. & Frangogiannis, N. G. The role of Smad signaling cascades in cardiac fibrosis. Cell Signal 77, 109826 (2021).

62. Zhang, M. et al. Notch3 Ameliorates Cardiac Fibrosis After Myocardial Infarction by Inhibiting the TGF-β1/Smad3 Pathway. Cardiovasc Toxicol 16, 316–324 (2016).

63. Lu, Q. et al. Targeting peroxisome proliferator-activated receptors: A new strategy for the treatment of cardiac fibrosis. Pharmacol Ther 219, 107702 (2021).

64. Liu, H. J., Liao, H. H., Yang, Z. & Tang, Q. Z. Peroxisome proliferator-activated receptor-γ is critical to cardiac fibrosis. PPAR Res 2016, (2016).

65. Li, X. et al. Aspirin Reduces Cardiac Interstitial Fibrosis by Inhibiting Erk1/2-Serpine2 and P-Akt Signalling Pathways. Cellular Physiology and Biochemistry 45, 1955–1965 (2018).

66. Zhang, Z., Yang, Z., Wang, S., Wang, X. & Mao, J. Targeting MAPK-ERK/JNK pathway: A potential intervention mechanism of myocardial fibrosis in heart failure. Biomedicine & Pharmacotherapy 173, 116413 (2024).

67. Mohamad, H. E., Askar, M. E. & Hafez, M. M. Management of cardiac fibrosis in diabetic rats; The role of peroxisome proliferator activated receptor gamma (PPAR-gamma) and calcium channel blockers (CCBs). Diabetol Metab Syndr 3, 1–12 (2011).

68. Pillarisetti, K. & Gupta, S. K. Cloning and relative expression analysis of rat stromal cell derived factor-1 (SDF-1)1: SDF-1 α mRNA is selectively induced in rat model of myocardial infarction. Inflammation 25, 293–300 (2001).

69. Boyle, A. J. et al. Myocardial production and release of MCP-1 and SDF-1 following myocardial infarction: Differences between mice and man. J Transl Med 9, 1–8 (2011).

70. Chen, Y. et al. Testosterone replacement therapy promotes angiogenesis after acute myocardial infarction by enhancing expression of cytokines HIF-1a, SDF-1a and VEGF. Eur J Pharmacol 684, 116–124 (2012).

71. Zhong, J. & Rajagopalan, S. Dipeptidyl peptidase-4 regulation of SDF-1/CXCR4 axis: Implications for cardiovascular disease. Front Immunol 6, 159972 (2015).

72. Döring, Y., Pawig, L., Weber, C. & Noels, H. The CXCL12/CXCR4 chemokine ligand/receptor axis in cardiovascular disease. Front Physiol 5 JUN, 88349 (2014).

73. Wen, J., Zhang, J.-Q., Huang, W. & Wang, Y. SDF-1α and CXCR4 as therapeutic targets in cardiovascular disease. Am J Cardiovasc Dis 2, 20 (2012).

74. Fiordelisi, A., Iaccarino, G., Morisco, C., Coscioni, E. & Sorriento, D. NFkappaB is a Key Player in the Crosstalk between Inflammation and Cardiovascular Diseases. International Journal of Molecular Sciences 2019, Vol. 20, Page 1599 20, 1599 (2019).

75. Cheng, W. et al. NF-κB, A Potential Therapeutic Target in Cardiovascular Diseases. Cardiovascular Drugs and Therapy 2022 37:3 37, 571–584 (2022).

76. Przybyszewski, E. M., Targher, G., Roden, M. & Corey, K. E. Nonalcoholic Fatty Liver Disease and Cardiovascular Disease. Clin Liver Dis (Hoboken) 17, 19–22 (2021).

77. Abdullah, M., Berthiaume, J. M. & Willis, M. S. Tumor necrosis factor receptor-associated factor 6 as a nuclear factor kappa B-modulating therapeutic target in cardiovascular diseases: at the heart of it all. Translational Research 195, 48–61 (2018).

78. Huang, W. et al. MicroRNA-3614 regulates inflammatory response via targeting TRAF6-mediated MAPKs and NF-κB signaling in the epicardial adipose tissue with coronary artery disease. Int J Cardiol 324, 152–164 (2021).

79. Ghigo, A., Laffargue, M., Li, M. & Hirsch, E. PI3K and Calcium Signaling in Cardiovascular Disease. Circ Res 121, 282–292 (2017).

80. Ghigo, A., Morello, F., Perino, A. & Hirsch, E. Therapeutic Applications of PI3K Inhibitors in Cardiovascular Diseases. Future Med Chem 5, 479–492 (2013).

81. Qin, W., Cao, L. & Massey, I. Y. Role of PI3K/Akt signaling pathway in cardiac fibrosis. Molecular and Cellular Biochemistry *2021* 476:*11* **476**, 4045–4059 (2021).

82. Chong, Z. Z., Shang, Y. C. & Maiese, K. Cardiovascular Disease and mTOR Signaling. Trends Cardiovasc Med 21, 151–155 (2011).

83. Yang, Z. & Ming, X. F. mTOR signalling: the molecular interface connecting metabolic stress, aging and cardiovascular diseases. Obesity Reviews 13, 58–68 (2012).

84. Sciarretta, S., Forte, M., Frati, G. & Sadoshima, J. New Insights Into the Role of mTOR Signaling in the Cardiovascular System. Circ Res 122, 489–505 (2018).

85. Samidurai, A., Kukreja, R. C. & Das, A. Emerging role of mTOR signaling-related miRNAs in cardiovascular diseases. Oxid Med Cell Longev 2018, (2018).

86. Das, A. & Reis, F. mTOR Signaling: New Insights into Cancer, Cardiovascular Diseases, Diabetes and Aging. International Journal of Molecular Sciences 2023, Vol. 24, Page 13628 24, 13628 (2023).

87. Ghafouri-Fard, S. et al. Interplay between PI3K/AKT pathway and heart disorders. Mol Biol Rep 49, 9767–9781 (2022).

88. Shen, A. R. et al. Integrin, Exosome and Kidney Disease. Front Physiol 11, 627800 (2021).

89. van den Hoogen, P., et al. Increased circulating IgG levels, myocardial immune cells and IgG deposits support a role for an immune response in pre- and end-stage heart failure. J Cell Mol Med 23, 7505–7516 (2019).

90. Tsimikas, S. et al. Relationship of IgG and IgM autoantibodies to oxidized low density lipoprotein with coronary artery disease and cardiovascular events. J Lipid Res 48, 425–433 (2007).

91. Ronan, G., Bahcecioglu, G., Yang, J. & Zorlutuna, P. Cardiac tissue-resident vesicles differentially modulate anti-fibrotic phenotype by age and sex through synergistic miRNA effects. Biomaterials 311, (2024).

92. Chen, Y. & Wang, X. miRDB: an online database for prediction of functional microRNA targets. Nucleic Acids Res 48, D127 (2019).

93. Huang, H. Y. et al. miRTarBase update 2022: an informative resource for experimentally validated miRNA–target interactions. Nucleic Acids Res 50, D222 (2021).

94. Agarwal, V., Bell, G. W., Nam, J. W. & Bartel, D. P. Predicting effective microRNA target sites in mammalian mRNAs. Elife 4, e05005 (2015).

95. Ashburner, M. et al. Gene ontology: tool for the unification of biology. The Gene Ontology Consortium. Nat Genet 25, 25–29 (2000).

96. Consortium, G. O. The Gene Ontology resource: enriching a GOld mine. Nucleic Acids Res 49, D325–D334 (2021).

97. Mi, H., Muruganujan, A., Ebert, D., Huang, X. & Thomas, P. D. PANTHER version 14: more genomes, a new PANTHER GO-slim and improvements in enrichment analysis tools. Nucleic Acids Res 47, D419–D426 (2019).

98. Liebermeister, W. et al. Visual account of protein investment in cellular functions. Proc Natl Acad Sci U S A 111, 8488–8493 (2014).

99. Tibshirani, R. Regression Shrinkage and Selection Via the Lasso. J R Stat Soc Series B Stat Methodol 58, 267–288 (1996).

100. Fan, J. & Li, R. Variable Selection via Nonconcave Penalized Likelihood and its Oracle Properties. J Am Stat Assoc 96, 1348–1360 (2001).

101. van Hoogevest, P. & Wendel, A. The use of natural and synthetic phospholipids as pharmaceutical excipients. European Journal of Lipid Science and Technology 116, 1088 (2014).

102. Le, N. T. T. et al. Soy Lecithin-Derived Liposomal Delivery Systems: Surface Modification and Current Applications. Int J Mol Sci 20, 4706 (2019).

103. Wang, G. & Wang, T. Oxidative stability of egg and soy lecithin as affected by transition metal ions and pH in emulsion. J Agric Food Chem 56, 11424–11431 (2008).

104. Yanasarn, N., Sloat, B. R. & Cui, Z. Nanoparticles Engineered from Lecithin-in-Water Emulsions As A Potential Delivery System for Docetaxel. Int J Pharm 379, 174 (2009).

105. Cornejo, M. A. & Linz, T. H. Selective miRNA quantitation with high-temperature thermal gel electrophoresis. Anal Chim Acta 1275, 341605 (2023).

106. Guo, M., Wei, Y., Lee, H., Maia, J. & Morrison, E. One-step extrusion of concentrated lidocaine lipid nanocarrier (LNC) dispersions. Int J Pharm 589, 119817 (2020).

107. Basara, G., Gulberk Ozcebe, S., Ellis, B. W. & Zorlutuna, P. Tunable human myocardium derived decellularized extracellular matrix for 3d bioprinting and cardiac tissue engineering. Gels 7, (2021).

