## Supplemental Data for "Comprehensive Multi-omics Mapping of Vesicle Cargo from Plasma or Novel Tissue Vesicles Reveals Pathological Changes in Cargo with Patient Age"

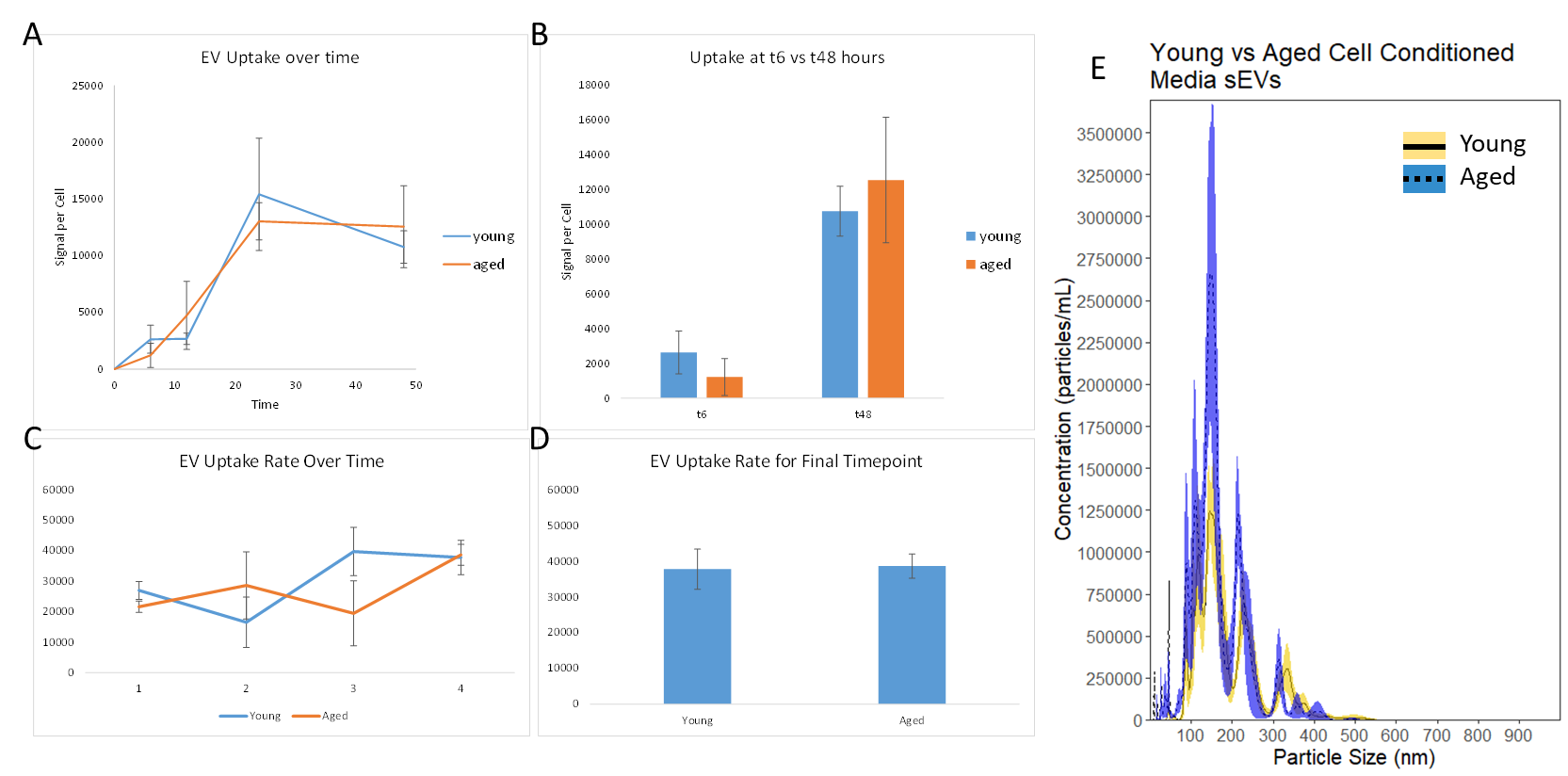

**Supplemental Figure S1: *Vesicle production and uptake assays with differently aged fibroblast cell lines.*** Cell uptake assays were performed with sEVs isolated from human hearts on young (P4) or aged (P14, artificially aged) human iCFs. Both for uptake over time (A, B) and uptake rate over time (C, D), no significant differences were observed at any timepoint. Additionally, sEVs gathered from media conditions by these cells did not demonstrate the observed difference between young and aged heart tissue sEVs. Statistics were performed via one-way ANOVA for A-D. * p < 0.05.

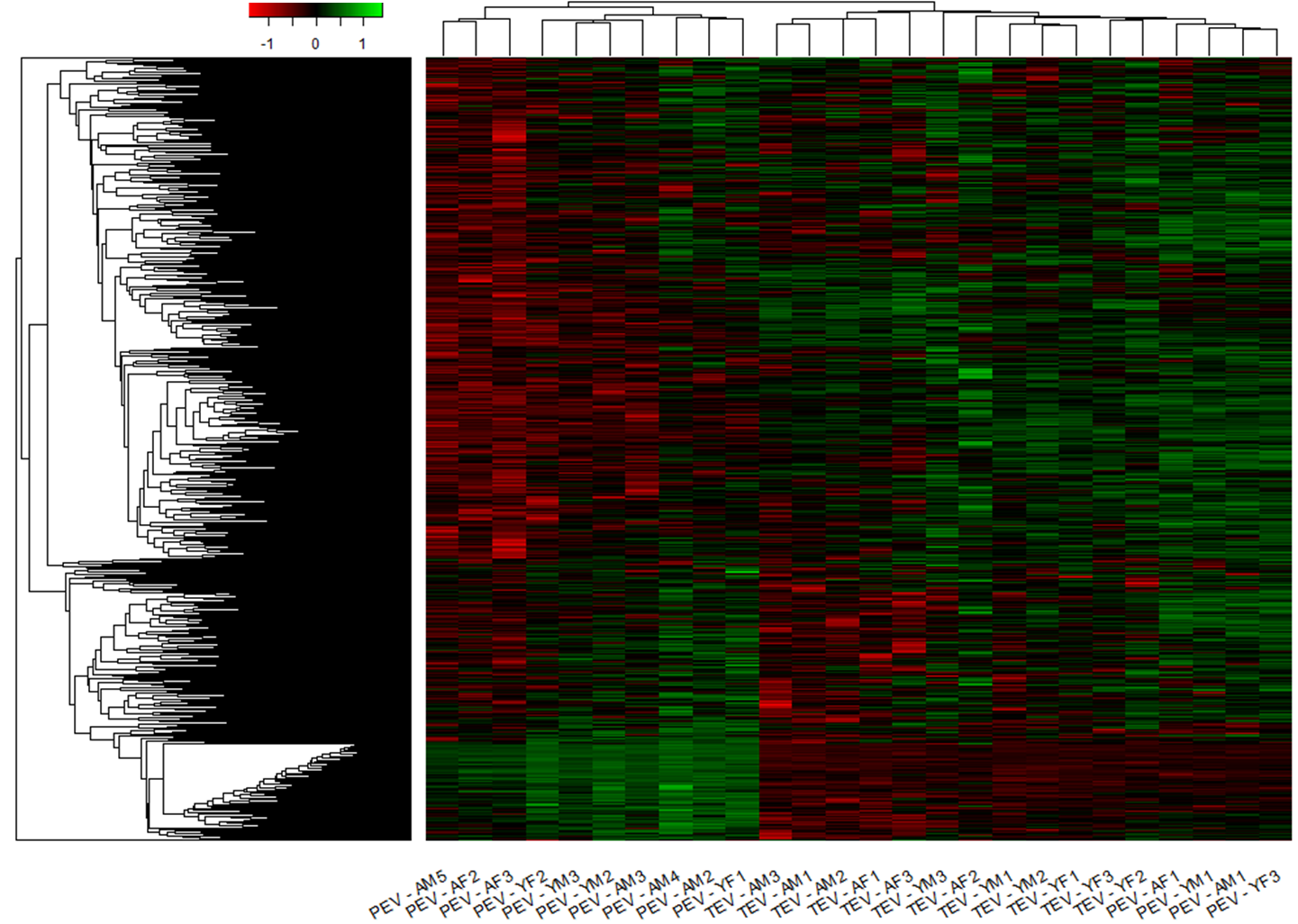

**Supplemental Figure S2: *Heatmap showing total miRNA profile for individual biological replicates of each cohort*.** Heatmap of the miRNA profiling results for each individual replicate of TEVs and PEVs. Clustering was performed in R using Euclidian distance calculations.

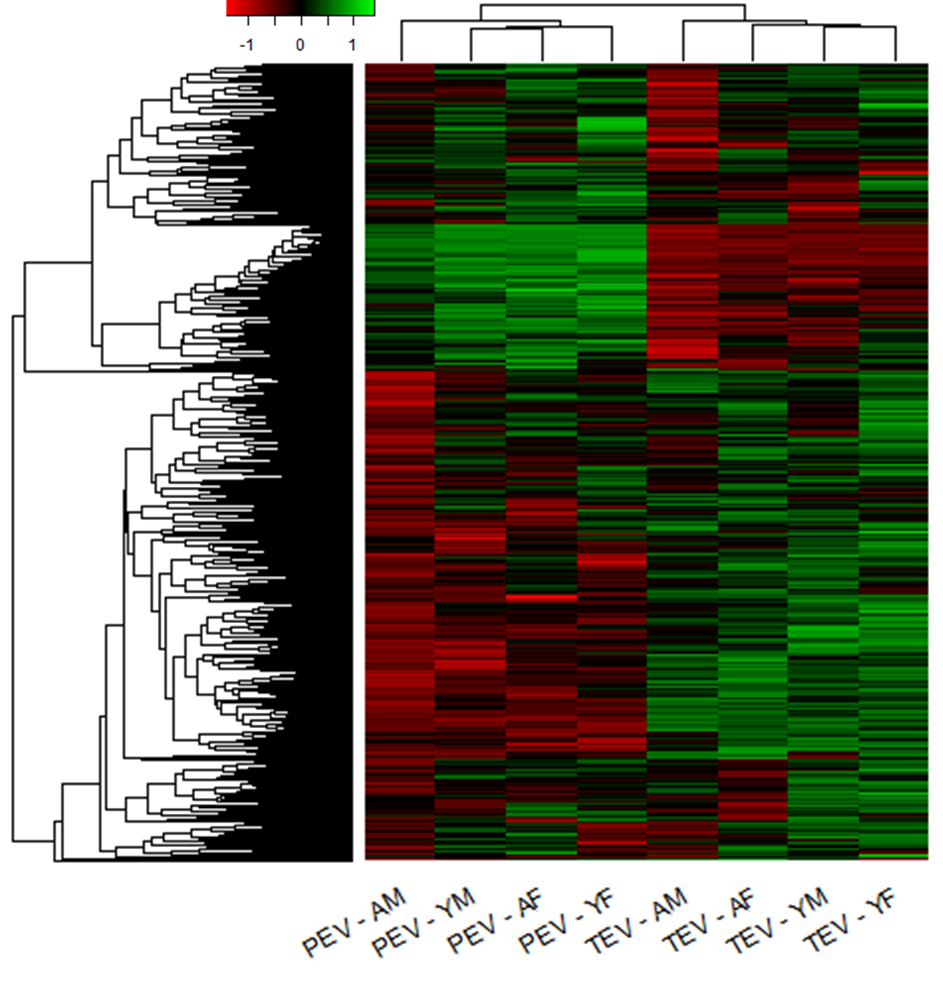

**Supplemental Figure S3: *Heatmaps showing total miRNA profile for aged and young TEV and PEV cohorts also separated by sex*.** Heatmap of the miRNA profiling data for TEVs and PEVs with cohorts grouped according to both age and sex. A and Y represent Aged or Young, and M and F represent Male and Female. Clustering was performed in R using Euclidian distance calculations.

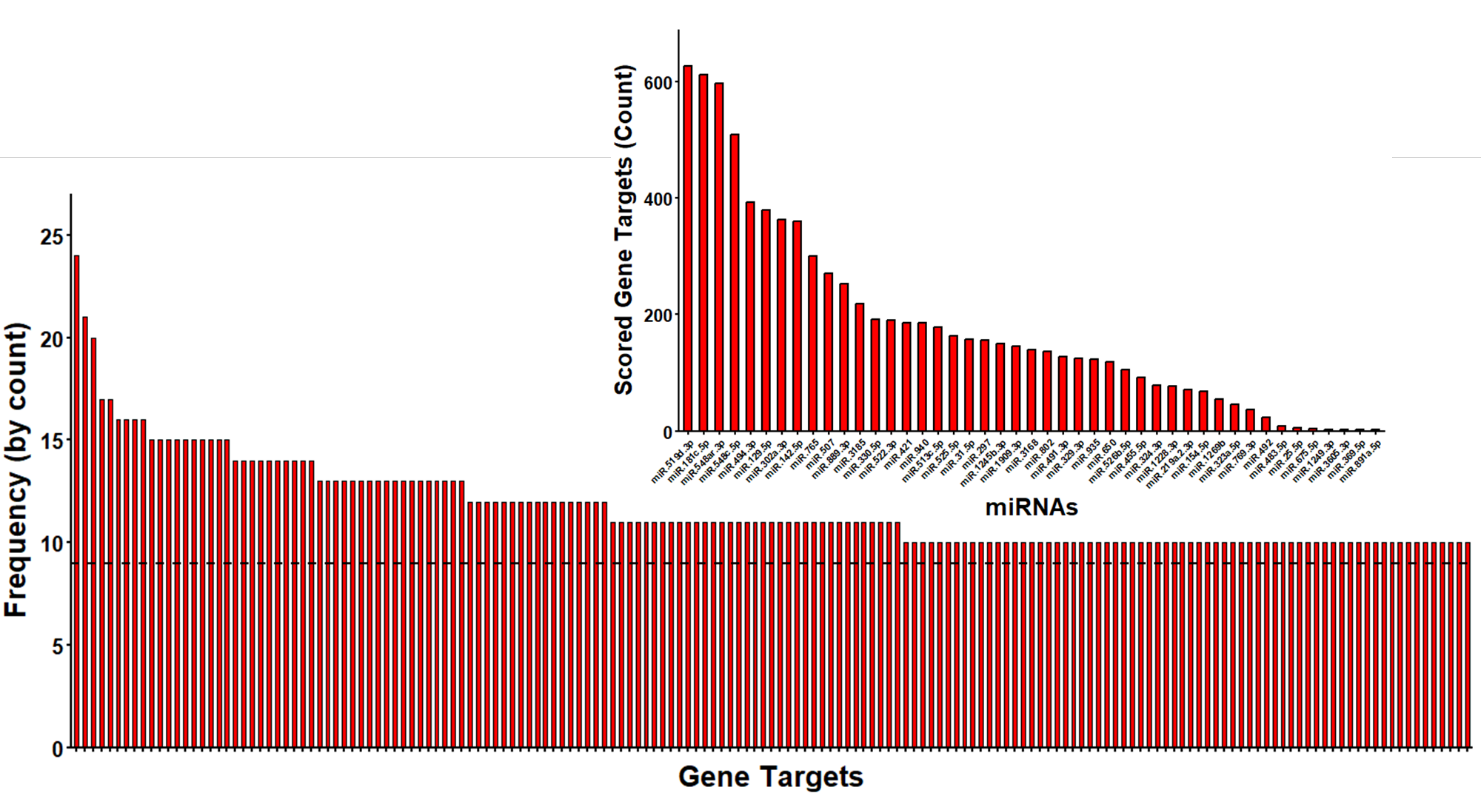

**Supplemental Figure S4: *Contribution of miRNAs to predicted gene targets and predicted gene targets selected for analysis*.** Histograms showing the predicted gene targets for the 45 target miRNAs (below), as well as the number of genes that each target miRNA is contributing to (above). Gene targets were only considered for analysis and included in the graph if identified as a potential target for at least 10 of the 45 target miRNAs, with the cutoff line (9 and below) indicated on the graph. A total list of gene targets can be found in Supplemental Table S6.

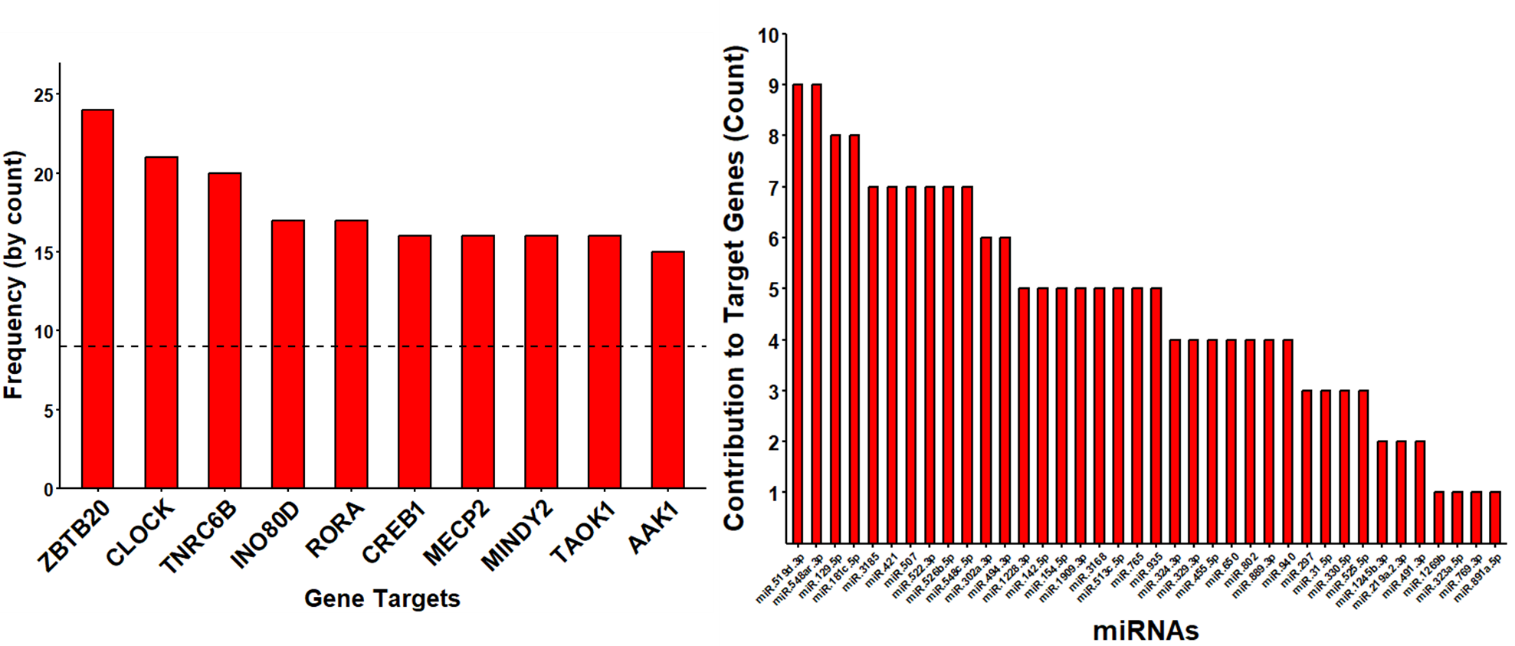

**Supplemental Figure S5: *10 most predicted gene targets of the target miRNAs and corresponding miRNA contribution*.** Histograms showing the top 10 most targeted genes of the 167 potential gene targets (left) and the contributing miRNAs were counted (right). Interestingly, no single miRNA contributed to all 10 gene targets.

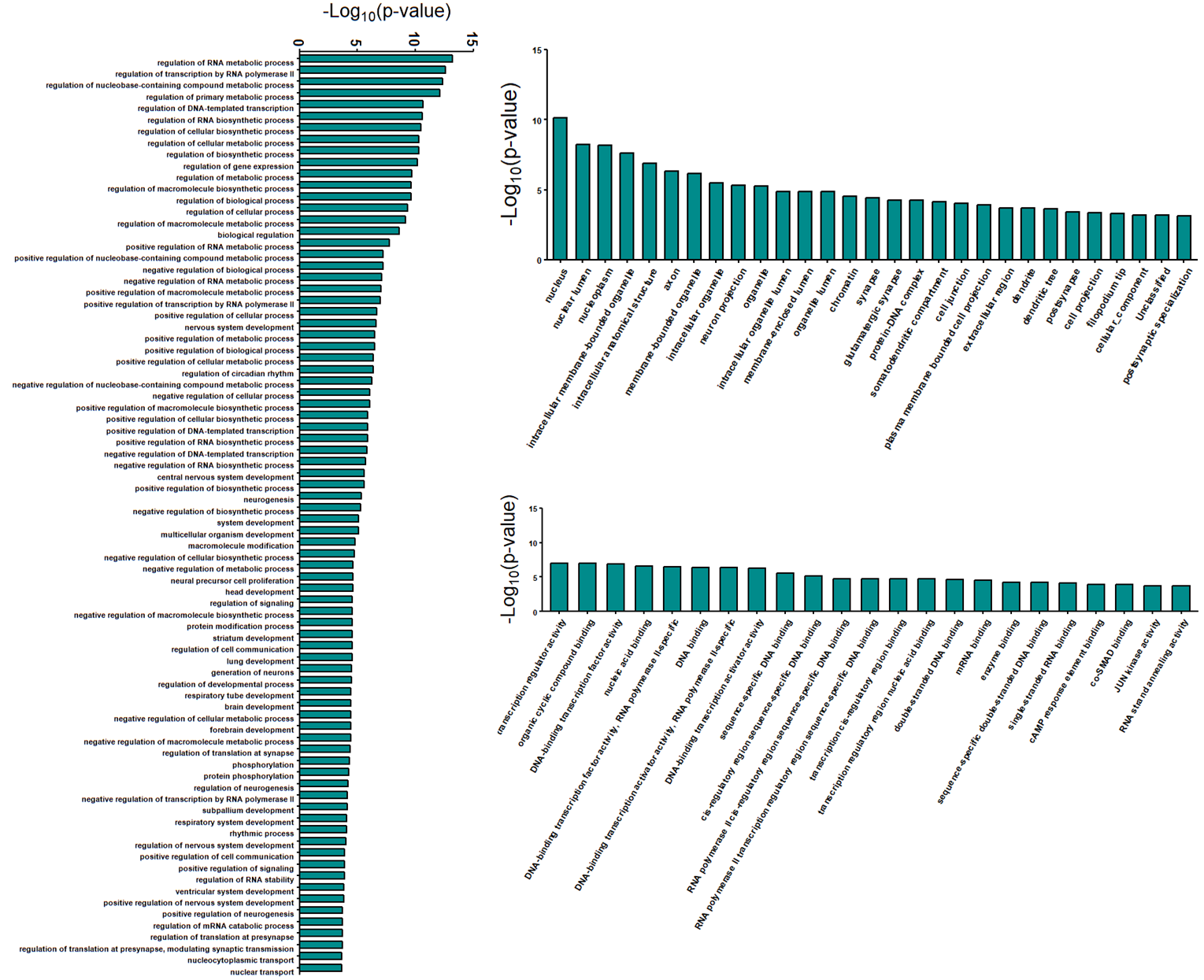

**Supplemental Figure S6: *Full gene ontology analysis of the 167 predicted gene targets*.** Full graphs for the gene ontology analysis of the identified 167 predicted gene targets for Biological Function (left), Cellular Component (top), and Molecular Function (bottom).

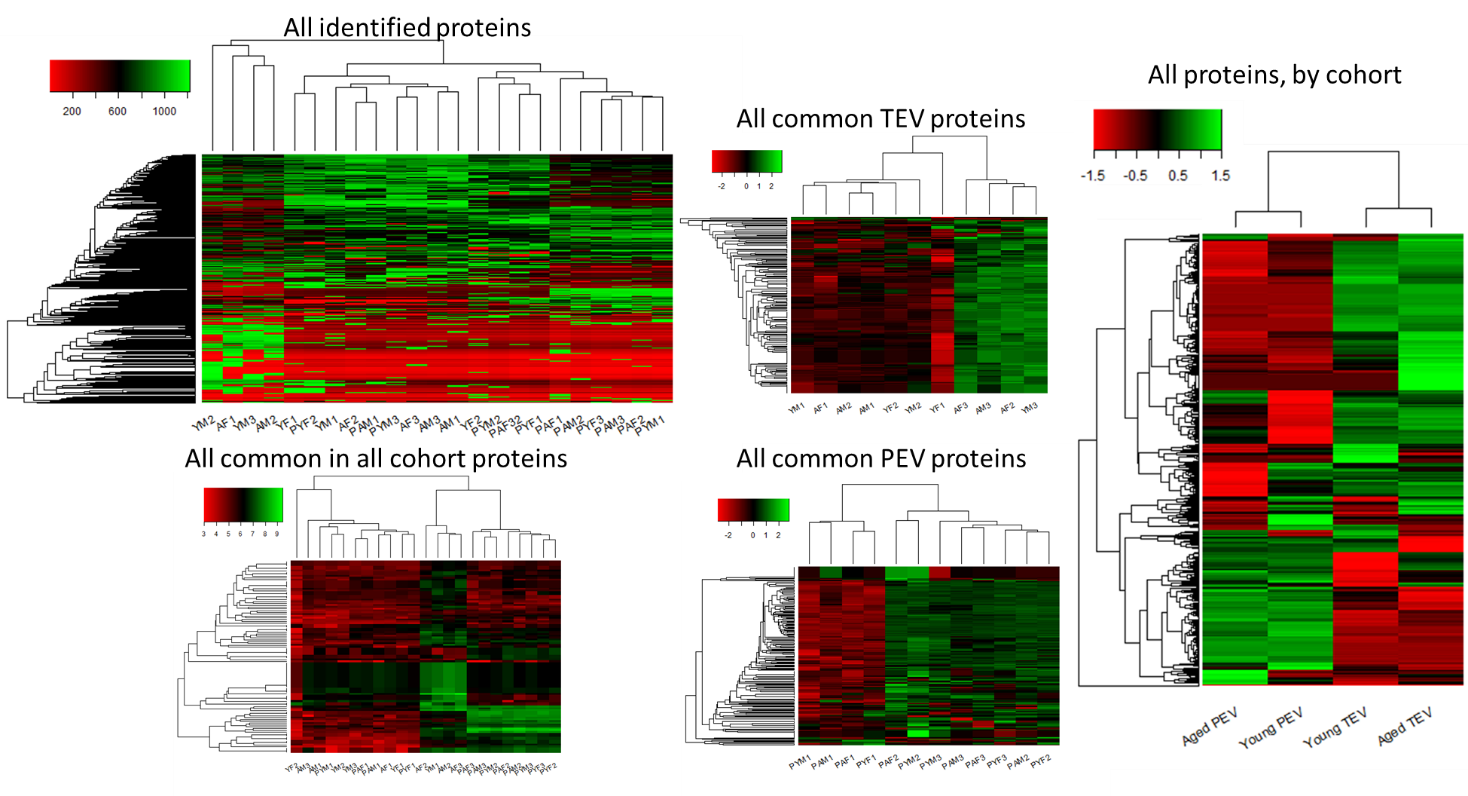

**Supplemental Figure S7: *Heatmaps for individual biological replicates of all identified proteins and for all proteins common to all cohorts*.** Heatmaps for the mass spectrometry proteomics results, divided by cohort. Total results, including null values, were mapped by cohort before subdivision (far right). All proteins by frequency of identification for all biological replicates (top left) was simplified into a heatmap containing proteins which were common in every biological replicate of both TEVs and PEVs for all cohorts and Z-scored (bottom left). Protein were also subdivided among those which were common in all TEV biological replicates (top right) and all PEV biological replicates (bottom right), each Z-scored respectively. P indicates a plasma sample, where no P indicates a tissue sample. A and Y stand for aged and young respectively, and M and F stand for male and female respectively.

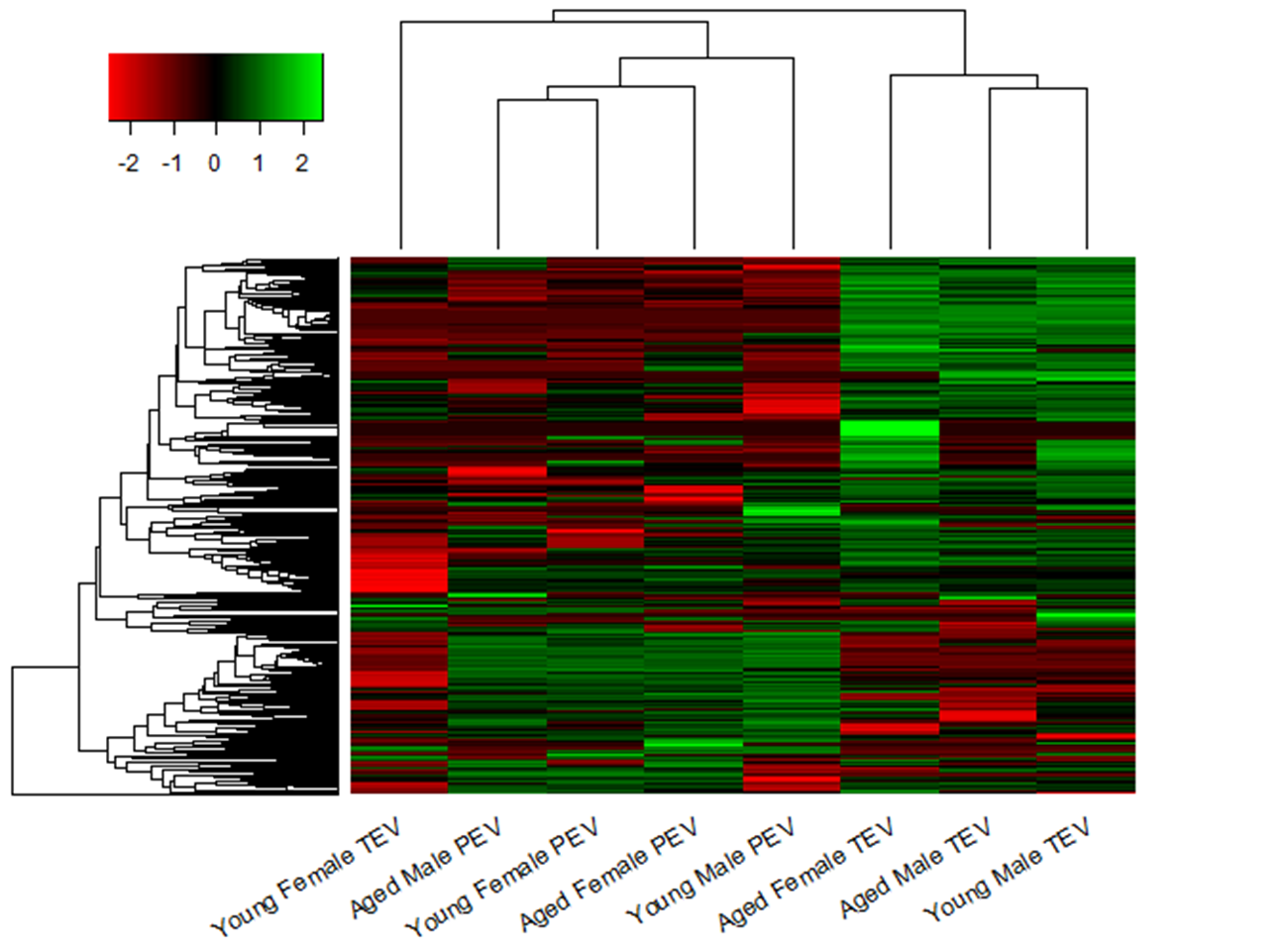

**Supplemental Figure S8: *Heatmaps of all identified proteins and for all proteins common to all cohorts grouped by both age and sex*.** Heatmap of the proteins common to all biological replicates of all cohorts, grouped by both age and sex for TEVs and PEVs.

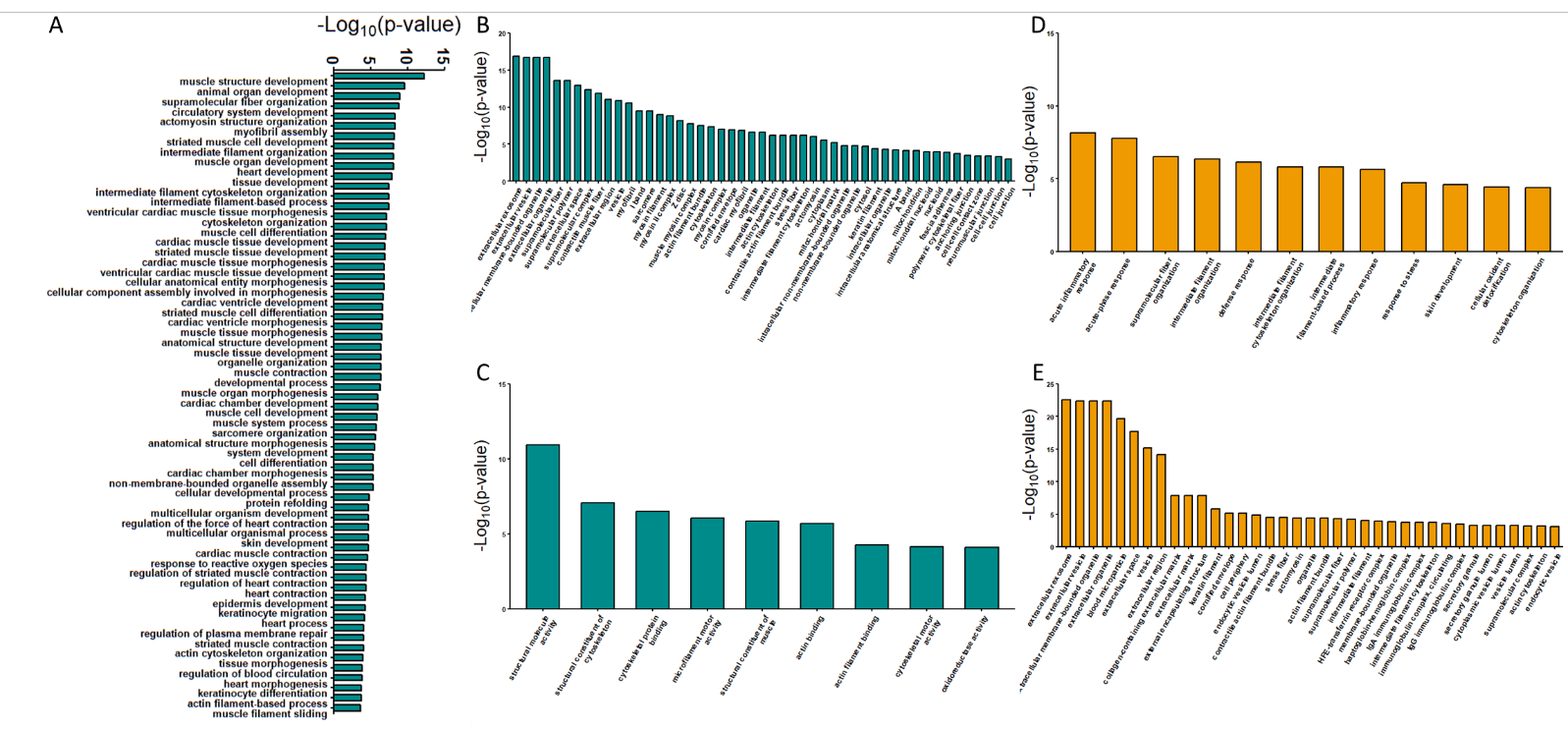

**Supplemental Figure S9: *Full gene ontology analysis of enriched proteins in TEVs and PEVs*.** Full graphs for the gene ontology analysis of the unique proteins for TEVs (cyan, A-C) and PEVs (orange, D-E). Data is shown for Biological Function (A, D), Cellular Component (B, E), and Molecular Function (C). No molecular function data is shown for PEVs because no statistically significant pathways were identified.

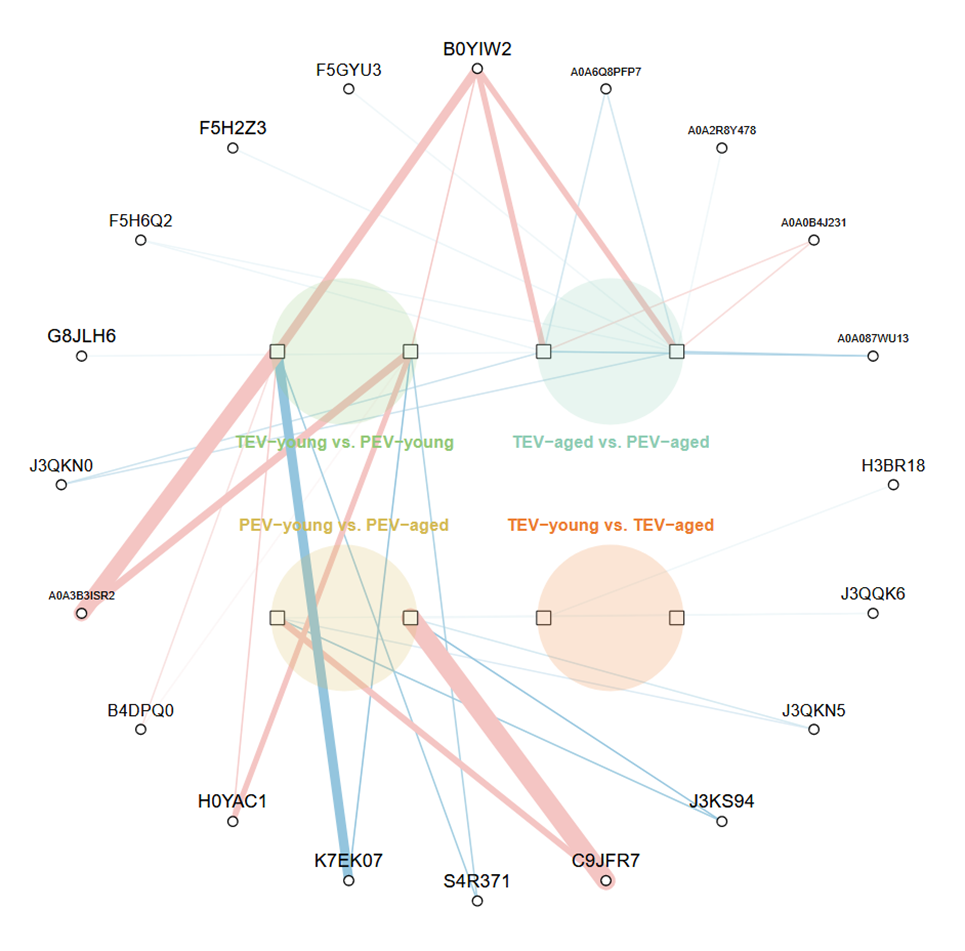

**Supplemental Figure S10: *Network map from machine learning analysis of protein dataset*.** A network plot representing the biomarkers selected from eight regularized logistic regression in 4 pairs of cohort comparisons: TEV-young vs. PEV-young; TEV-aged vs. PEV-aged; TEV-young vs. TEV-aged; PEV-young vs. PEV-aged, each depicted by a colored circle at the center of the network. The square nodes represent the eight models (with LASSO and SCAD penalties in one full dataset and three biologically defined subsets); the circular nodes along the periphery represent individual miRNA selected by at least one model. Edges connect miRNAs to the models in which they were selected, with red edges indicating a positive coefficient and blue edges indicating a negative coefficient. The thickness of each edge reflects the absolute magnitude of the coefficient.

**Supplemental Table S1: *Donor Information***

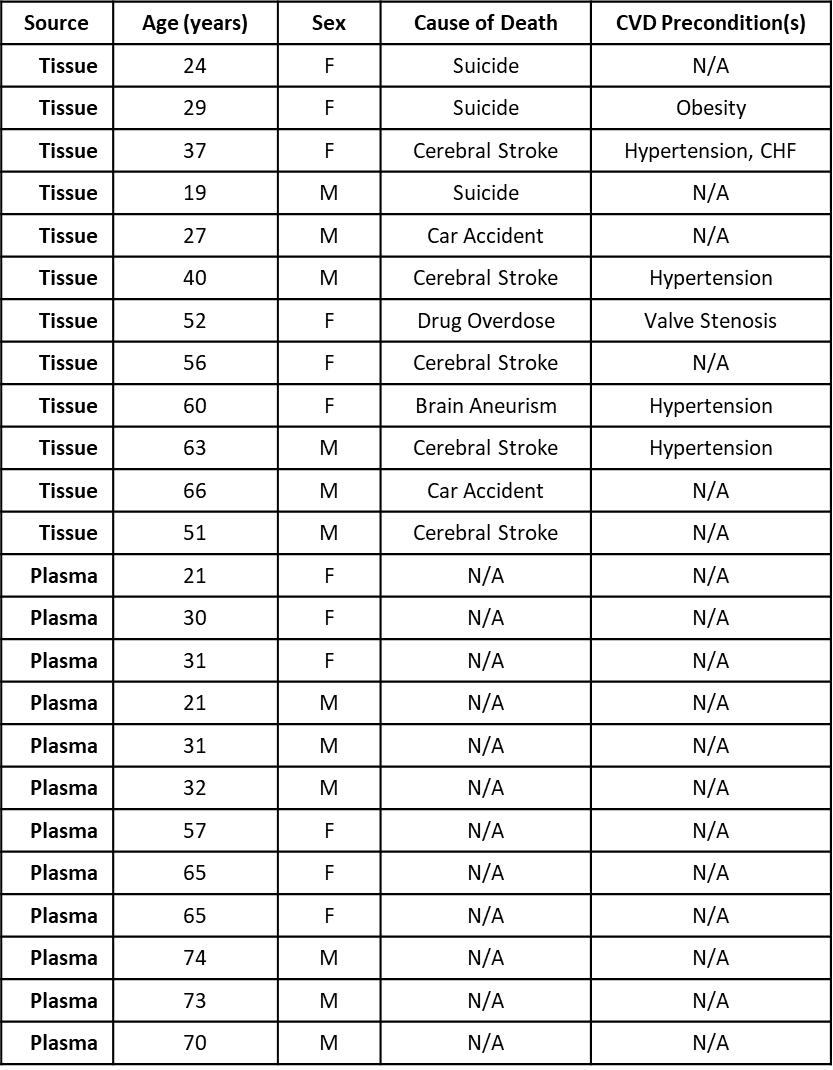

**Supplemental Table S2: *Nanoparticle Tracking Analysis Statistics***

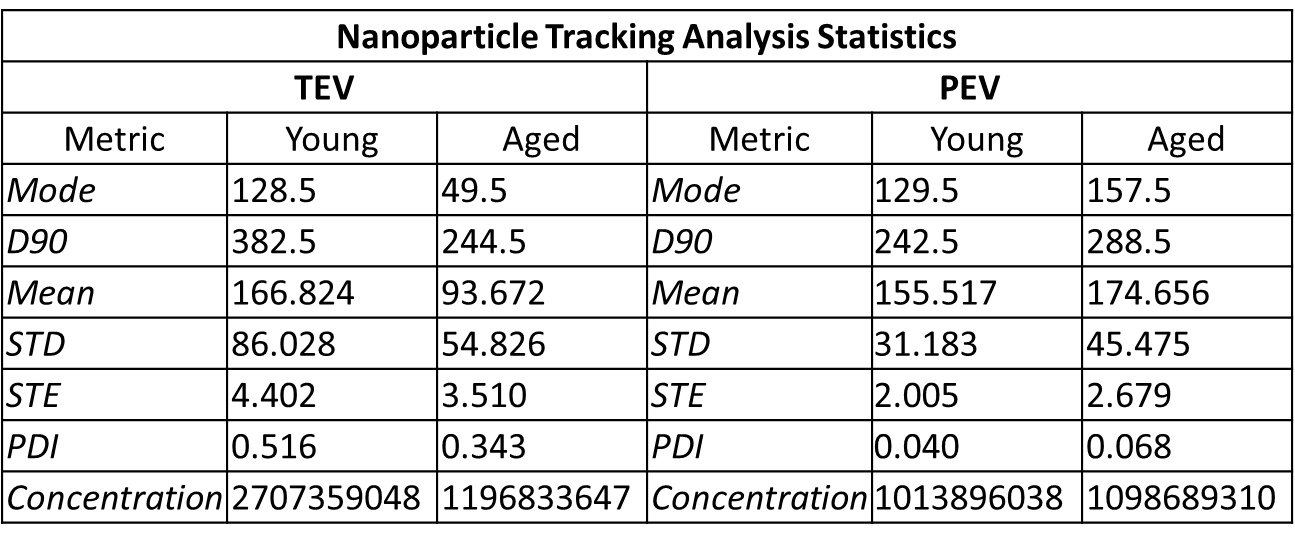

**Supplemental Table S3: *Relative Expression Values for all Detected miRNAs in TEVs and PEVs***

Please see attached file: Supplemental Table S3

**Supplemental Table S4: *Differentially Abundant miRNAs Enriched Uniquely in TEVs or PEVs***

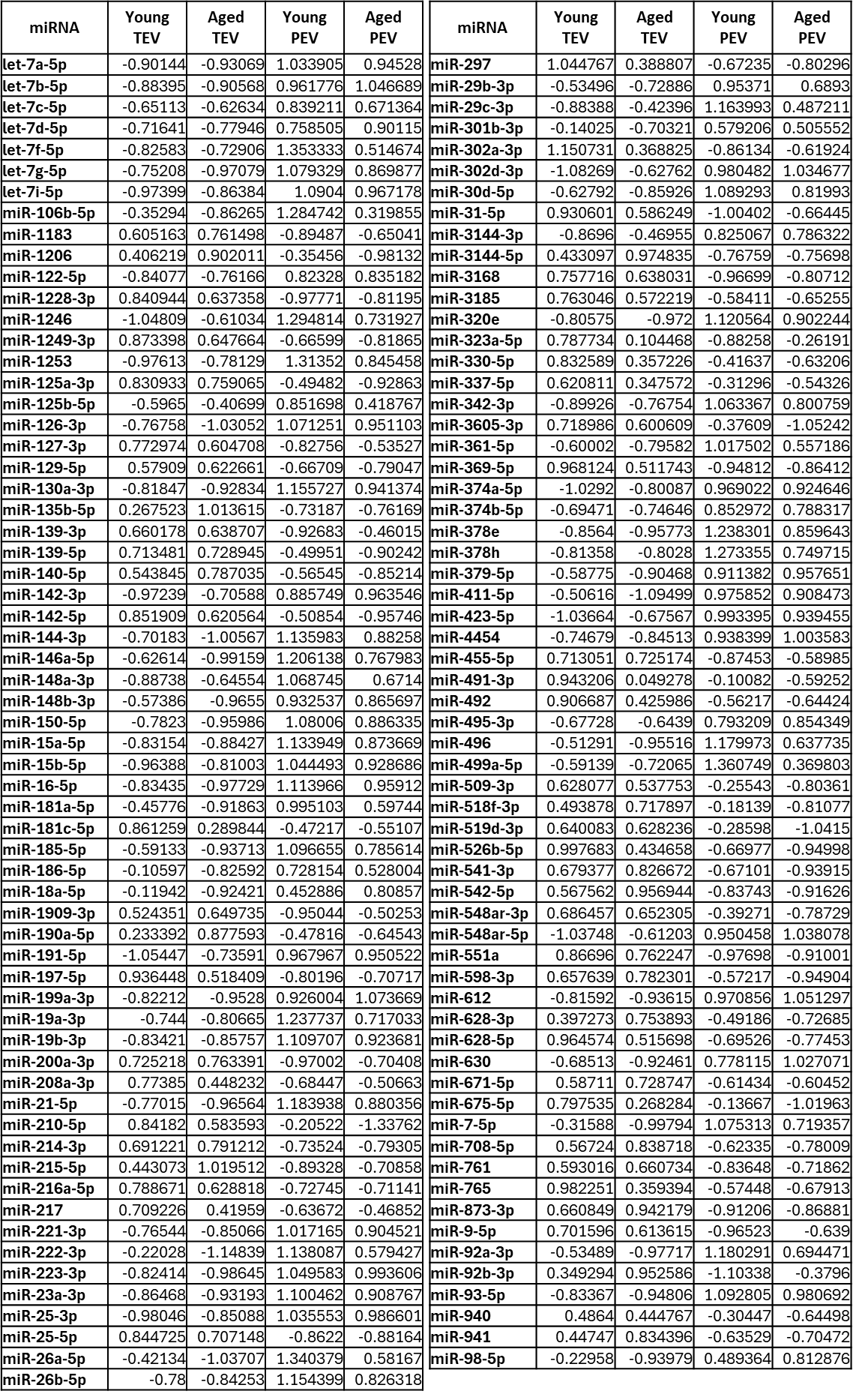

**Supplemental Table S5: *List of predicted gene targets for each target miRNA***

Please see attached file: Supplemental Table S5

**Supplemental Table S6: *167 Gene Targets Selected for Analysis and Corresponding miRNA Contributors***

Please see attached file: Supplemental Table S6

**Supplemental Table S7: *Relative Expression of all Common Proteins in TEVs and PEVs***

Please see attached file: Supplemental Table S7

**Supplemental Table S8: *Proteins Uniquely Expressed in TEVs or PEVs Overall***

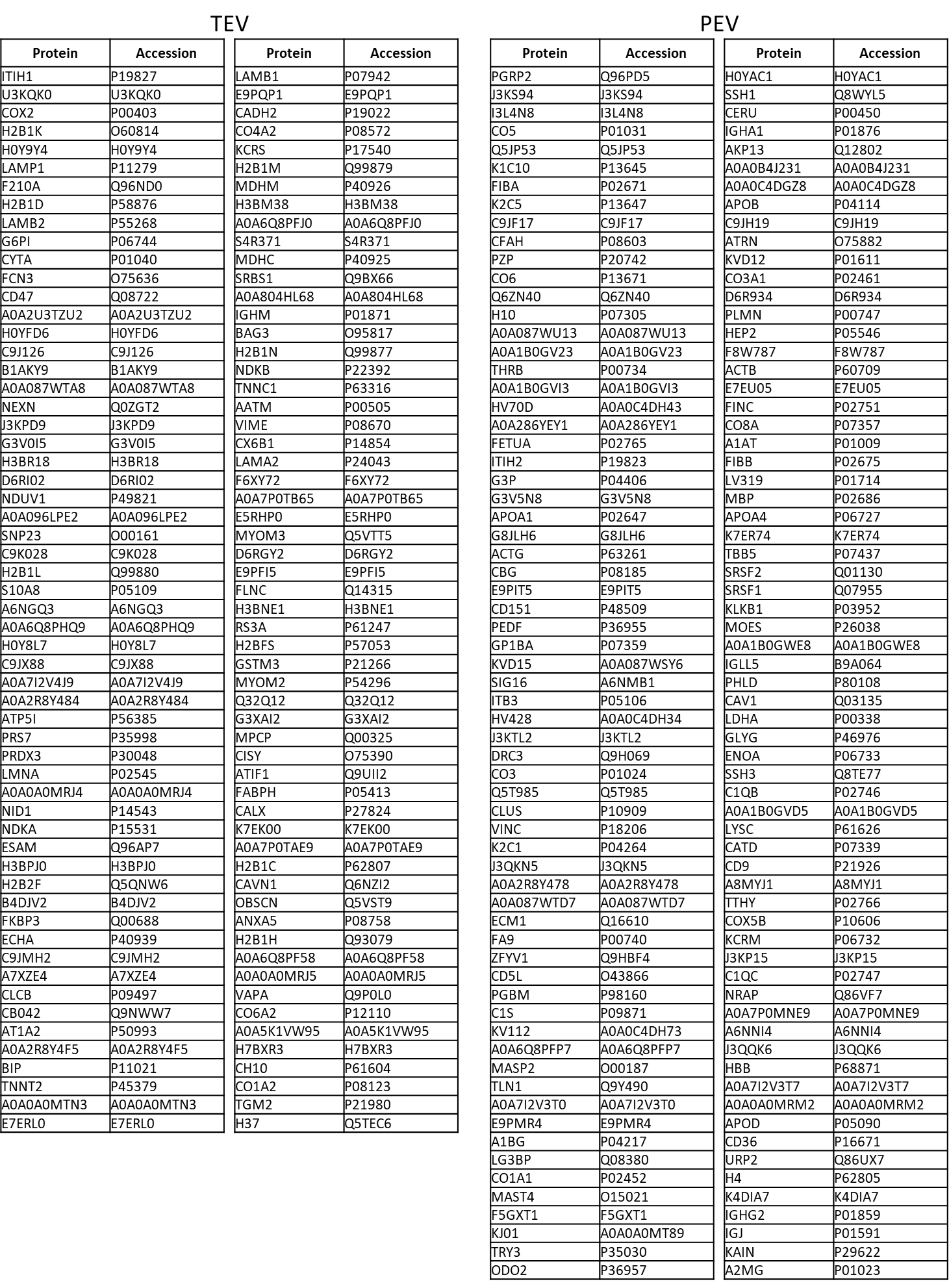

**Supplemental Table S9: *Proteins Uniquely Expressed in Aged TEVs or aged PEVs***

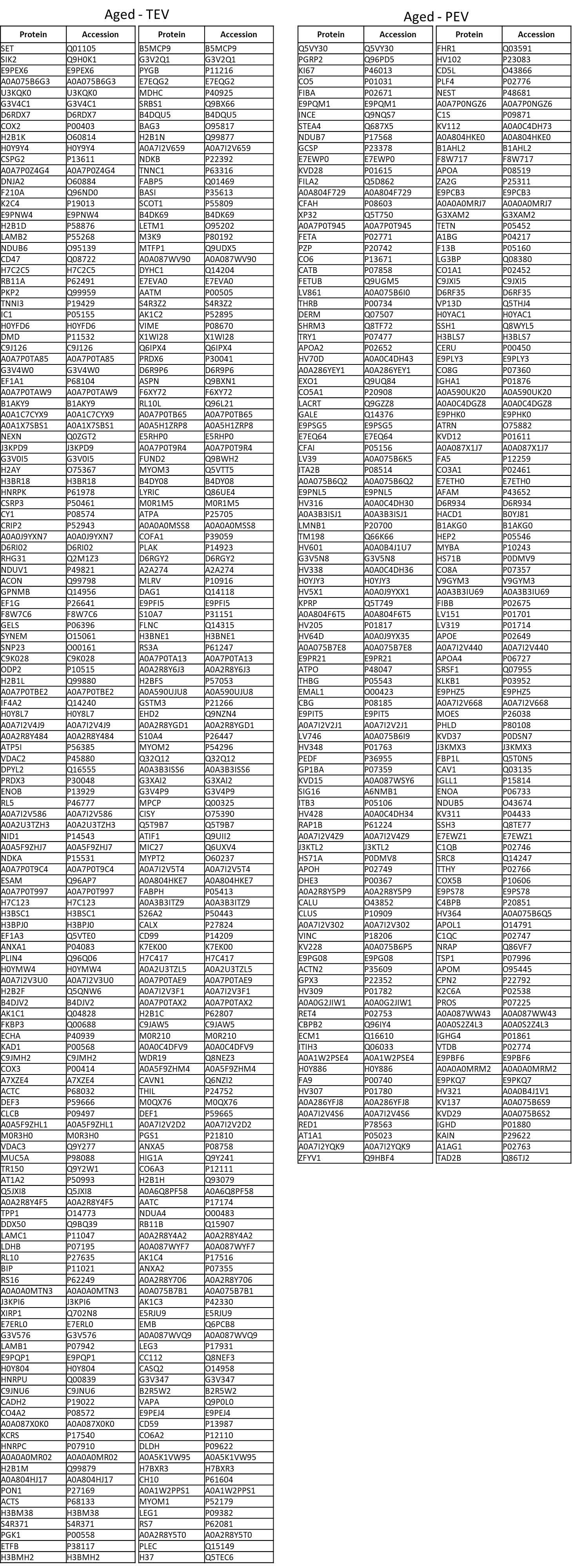

**Supplemental Table S10: *Machine learning-identified distinguishing biomolecules from profiling data***

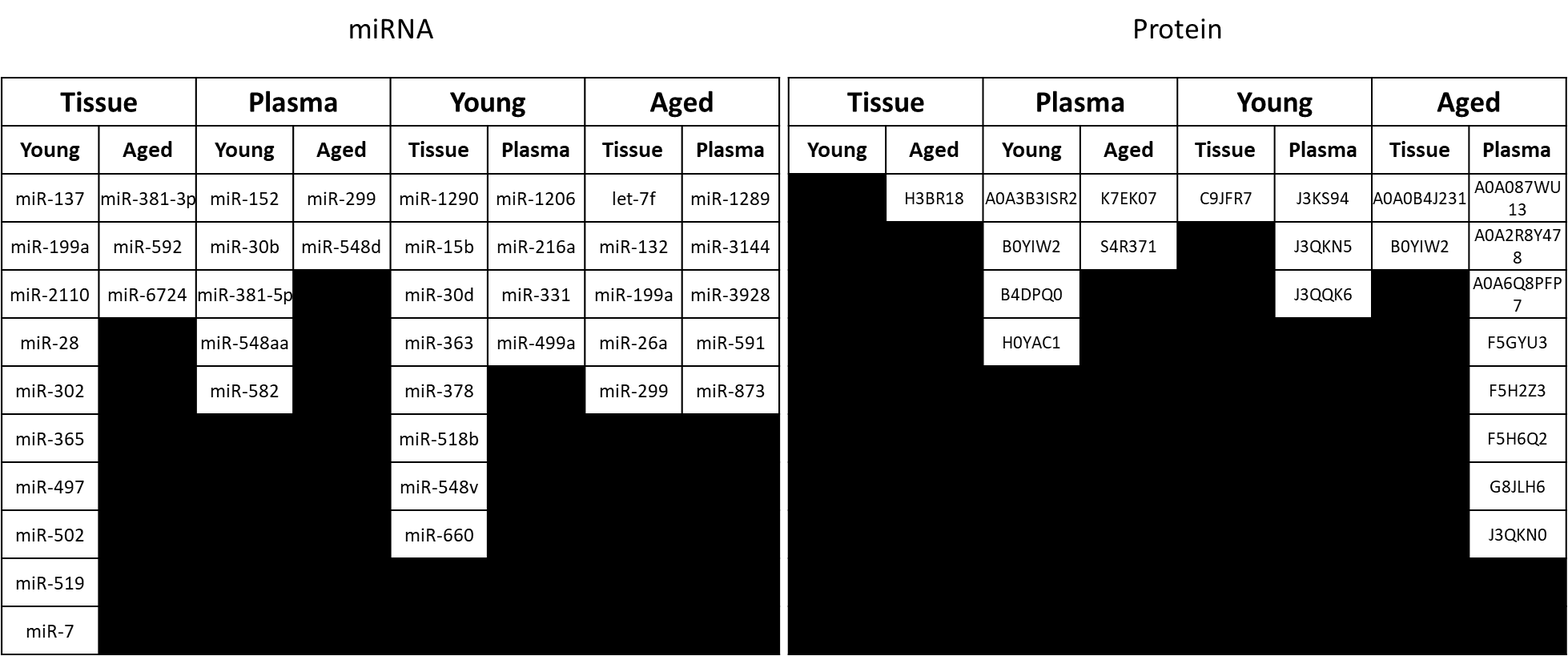
